# Absolute quantification and single-cell dose-response of cytosolic siRNA delivery

**DOI:** 10.1101/2021.04.21.440807

**Authors:** Hampus Hedlund, Hampus Du Rietz, Johanna Johansson, Wahed Zedan, Linfeng Huang, Jonas Wallin, Anders Wittrup

## Abstract

Endosomal escape and subsequent cytosolic delivery of small inhibitory RNA (siRNA) therapeutics is believed to be highly inefficient. Since, it has not been possible to quantify cytosolic amounts of delivered siRNA at therapeutic doses, determination of delivery bottlenecks and total efficiency has been difficult. Here, we present a confocal microscopy-based method to detect cytosolic delivery of fluorescently labelled siRNA during lipid-mediated delivery. This method enables detection and quantification of sub-nanomolar cytosolic siRNA release amounts from individual release events with measures of quantitation confidence for each event. Single-cell kinetics of siRNA-mediated knockdown in cells expressing destabilized eGFP unveiled a dose-response relationship with respect to knockdown induction, depth and duration in the range from several hundred to thousands of cytosolic siRNA molecules. Accurate quantification of cytosolic siRNA, and the establishment of the intracellular dose-response relationships, will aid the development and characterization of novel delivery strategies for nucleic acid therapeutics.

## Introduction

Small inhibitory RNA (siRNA) therapeutics are rapidly entering clinical use for multiple diseases. LNP formulated siRNA targeting transthyrein (patisiran)^1^ and three GalNAc-conjugated chemically stabilized free siRNA compounds (givosiran^2^, lumasiran^3^ and inclisiran^4^) have recently received clinical approval and several other substances are in clinical development. Both LNPs and GalNAc-conjugated siRNA target the liver, currently the organ most amenable to macromolecular delivery. A key impediment in efforts to improve siRNA delivery to other tissues has been a lack of tools to accurately detect and quantify successful cytosolic delivery of siRNA. Total tissue siRNA amounts do not directly correlate to biological effects due to inefficiency and variability of cellular internalization and endosomal escape of the delivered siRNA^5^. Crucially, methods to quantify the cytosolic concentration of a delivered siRNA have been lacking and the dose-response relationship for cytosol-delivered siRNA has not been clear. As a consequence, it has not been possible to determine delivery efficiencies, and the scope for improvement, for different delivery strategies.

Transfection lipid-mediated siRNA delivery has been shown to proceed by discrete release events resulting in a detectable cytosolic siRNA signal^6–8^. However, the detection level in previous experiments was not low enough to capture varying degrees of knockdown, instead all detected events resulted in maximal knockdown^8^. Other strategies to quantify absolute delivery amounts during lipid-mediated siRNA delivery, have relied on either electron-microscopy of fixed cells^9^ or ensemble measurements of Ago2-immunoprecipitated siRNA as a surrogate marker for cytoplasmic siRNA^10,11^. These non-live cell methods cannot be used to correlate single-cell delivery and knockdown and in addition, the accuracy and precision of the quantifications are difficult to evaluate. Intracellular dose-response determination of siRNAs has further been addressed using microinjection experiments with highly divergent results, suggesting cytosolic half-maximal inhibitory concentrations (IC50) or doses between 12 and several hundred siRNA molecules ^12–14^.

Here we present an imaging strategy in live cells to quantify endosomal escape events during lipid-mediated delivery with sufficiently few siRNA molecules to capture the dose-response interval with respect to knockdown outcomes in individual cells. Our method is based on array-confocal detection of fluorescent siRNA combined with post-acquisition processing to exclusively measure cytosolic (non-vesicular) siRNA. The method enables detection and absolute quantification of sub-nanomolar cytosolic siRNA release amounts from individual cytosolic release events and provides measures of quantitation confidence for each event. Single-cell kinetics of siRNA-mediated knockdown in cells expressing destabilized eGFP unveiled a dose-response relationship with respect to knockdown induction, depth and duration in the range from a few hundred to several thousand cytosolic siRNA molecules.

## Results

Our first aim was to improve the sensitivity and accuracy of cytosolic siRNA detection compared to previous efforts^8^. Ultimately, we wanted to detect and quantify release events in cells incubated with siRNA doses resulting in sub-maximal knockdown (that is, doses around typical IC50 values) for relatively efficient siRNA sequences. To this end, we set out to visualize transfection lipid-mediated endosomal escape in cells incubated with sub-nanomolar concentrations of Alexa Fluor 647-labelled siRNA (siRNA-AF647). Using a GaAsP array-confocal detector (Airyscan, ZEISS)^15^ frequent endosomal escape events and cytosolic dispersion of siRNA-AF647 was clearly visible during Lipofectamine 2000 transfection with 400 pM siRNA (Fig. 1a and Supplementary Movie 1), and also lower doses. In this imaging, we could take advantage of the high dynamic range of the array-confocal detector, stemming from the use of multiple small GaAsP-detectors that tolerate a high photon flux (in total, distributed over all detectors) while maintaining a very low detection limit and high rejection of out-of-focus light. Thus, an Airyscan array-confocal detector can be used to detect cytosolic dispersion of siRNA over a large field-of-view (FOV) during low-dose transfection.

**Figure 1.**
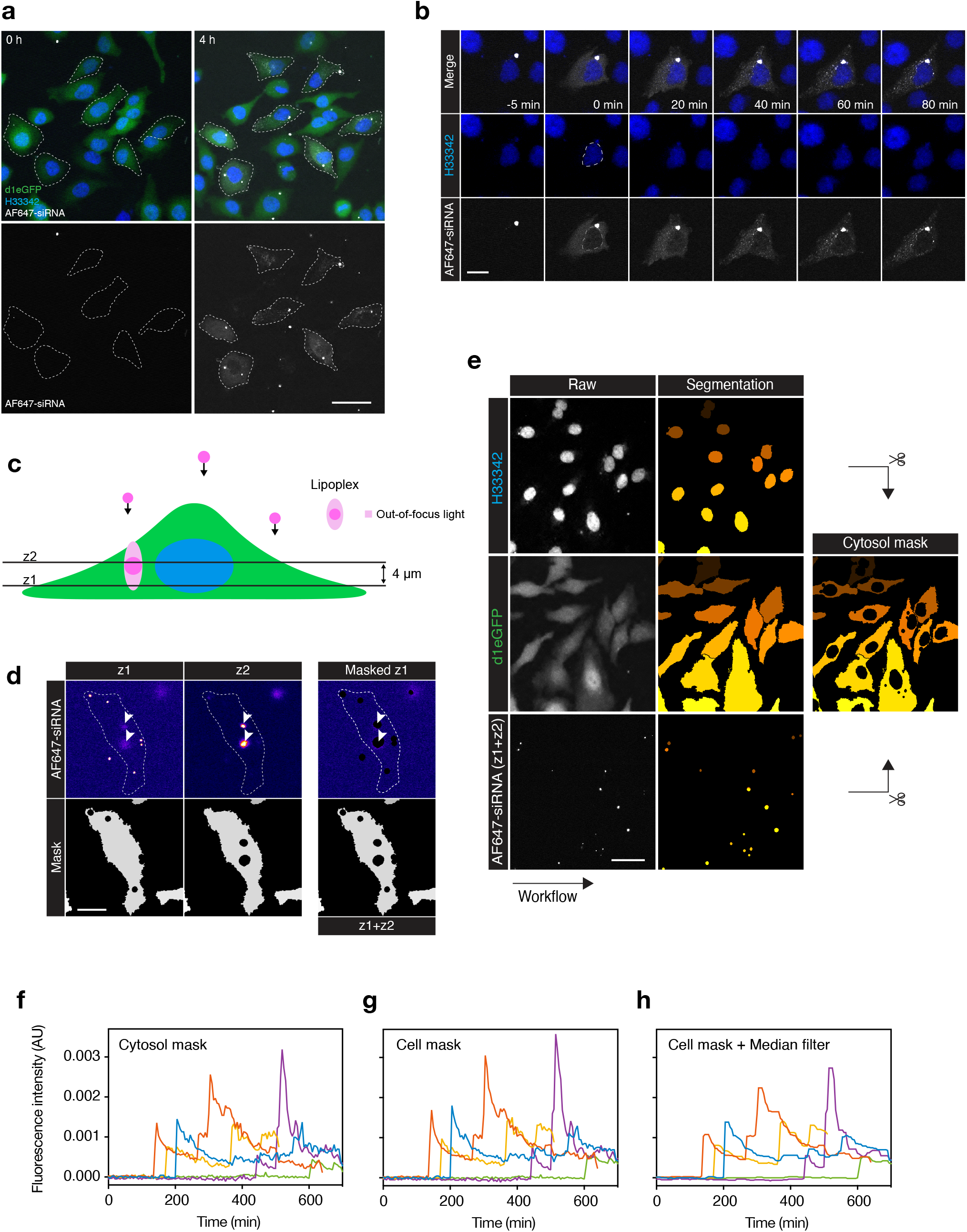
Detecting cytosolic delivery of siRNA during sub-nanomolar siRNA transfection. An Airyscan confocal detector was used to image HeLa cells stably expressing d1-eGFP every 5 min during treatment with lipoplexed siRNA-AF647. **(a)** Cells exhibiting cytosolic distribution of siRNA-AF647 after 4 h treatment with 0.4 nM siRNA. Scalebar is 50 μm. Representative images from 8 independent experiment. **(b)** Image detail showing a sudden endosomal release event and cytosolic dispersion of siRNA-AF647 at *t* = 0, during 0.2 nM siRNA transfection. Outline indicate nucleus. Scalebar is 20 μm. Representative images from 8 independent experiment. **(c)** Two *z-*planes are acquired during live-cell microscopy. In-focus lipoplexes captured in the upper plane (z_2_) is used to mask out-of-focus light contaminating the lower *z*-plane (z_1_) used for cytosolic siRNA detection. **(d)** Cell masked from bright lipoplexes, including out-of-focus light (visible in z_1_) caused by siRNA-AF647 lipoplexes at an elevated plane (z_2_). Outline indicate cell and arrows indicate lipoplex fluorescence detected in both *z*-planes. Scalebar is 20 μm. **(e)** The final cytosol image mask used for siRNA detection after segmentation and removing nuclei and lipoplexes. Scalebar is 50 μm. **(f–h)** The median siRNA-AF647 fluorescence intensity was measured in tracked cells (distinguished by color), **(f)** using a cytosol mask (nucleus and lipoplexes removed), **(g)** cell mask (lipoplexes removed), or **(h)** cell mask with a one-dimensional median filtering of the measurements using a 5-frame moving window. Cells were treated with 0.67 nM lipoplexed siRNA. Data is representative of 8 independent experiment. A constant siRNA-to-lipid ratio (2 pmol:4 μL) was used for all experiments.

Fluorescence intensities can be converted to absolute concentrations of a fluorescently labelled analyte through the use of reference measurements of samples with known concentrations. Our imaging setup had linear fluorescence readout for solutions containing 1 nM to 1000 nM siRNA-AF647 (R^2^ = 0.9998 ± 0.0002, mean ± s.d.) (Supplementary Fig. 1) Acquiring images at maximal FOV (beyond recommended settings) was associated with notable vignetting (Supplementary Fig. 2), which was later corrected during post-acquisition processing. For accurate (low-noise) quantification, cytosolic siRNA fluorescence should be measured in cross-sections of cells at the largest area possible, where the fluorescence intensity is homogenous (uniform cytosolic dispersion). Shortly after the release event, the cytosolic distribution of siRNA-AF647 is highly homogenous. However, 20–60 min after the release, the cytosolic siRNA progressively accumulates in cytoplasmic foci (Fig. 1b), as has been observed previously^6^. Consistently, siRNA was excluded from the nucleus upon cytosolic release during low-dose transfections. Thus, fluorescence measurements in the cytosol of cells within minutes after release, offers a potential to quantify the cytosolic siRNA concentration down to 1 nM and below.

To exclusively measure the cytosolic fraction of the siRNA-AF647 fluorescence, the contribution of non-cytosolic siRNA (non-released lipoplexes) had to be removed. While highly fluorescent lipoplex particles present in the focal plane can be easily delineated and masked from images based on their high signal intensity, the hazy fluorescence contribution of out-of-focus particles are not as easily excluded. To solve this, we acquired two confocal planes of cells incubated with lipoplexes, spaced 4 μm apart (Fig. 1c,d). The eGFP signal in the lower plane (*z*_1_) was used to identify boundaries of cells. Bright, in focus lipoplex particles present in either the lower (*z*_1_) or upper plane (*z*_2_) were identified and masked from both planes with an expanded margin in the siRNA-AF647 channel. In this way, the contribution of out-of-focus fluorescence from lipoplexes in the upper part of the cells could be excluded from the lower plane image. Finally, cell nuclei were masked in the siRNA channel, to obtain a homogenous cytosol devoid of fluorescence contribution from intact lipoplexes (Fig. 1e).

To facilitate unbiased analysis of cytosolic siRNA delivery in a large number of cells, we designed an algorithm to automatically detect cytosolic release events in time-lapse imaging datasets. Using the imaging strategy described above and cell tracking (Supplementary Fig. 3), we measured cellular siRNA fluorescence over time in individual cells, revealing sudden increases in the cytosolic fluorescence intensity of variable magnitude (Fig. 1f). Instances with abrupt siRNA signal increase were flagged as potential release events (Supplementary Fig. 4). To limit the number of false-positive events, the signal-to-noise ratio was improved by analyzing the entire cell including the nuclear region for release event detection, increasing the area of evaluation for a more robust event identification (Fig. 1g). Median filtering further decreased signal noise and made systematic intensity shifts more apparent (Fig. 1h). Thus, continuous monitoring of the cytosol enables automated detection of sudden siRNA release events using single-cell fluorescence intensity measurements.

To evaluate the performance of the event-calling algorithm and the sensitivity of the imaging setup, we used an independent assay based on the highly sensitive membrane damage sensor galectin-9^16^ (Fig 2a), shown to detect membrane damage associated with release of lipid-formulated or conjugated siRNA.^8,17^ Using live-cell microscopy of HeLa cells stably expressing YFP-tagged galectin-9, recruitment of galectin-9 to siRNA-AF647 lipoplexes upon endosomal escape served as ground truth for evaluating the detection of release events. In this way, we found that release events could be identified by the algorithm with 70% specificity and 76% sensitivity (Fig. 2b,c). Combined with manual control of the detected events, sensitivity and specificity was improved to 81% and 99%, respectively. In comparison, blinded manual inspection of time-lapse imaging datasets to identify sudden cytosolic dispersions of siRNA showed a near complete agreement with events detected by galectin-9 recruitment (92% sensitivity, 99% specificity), but this approach was very labor intensive. By combining the automated identification of release events with subsequent manual inspection, we achieved a workflow capable of detecting siRNA release with high specificity (no false events) and satisfactory sensitivity while keeping manual labor efforts reasonable.

**Figure 2.**
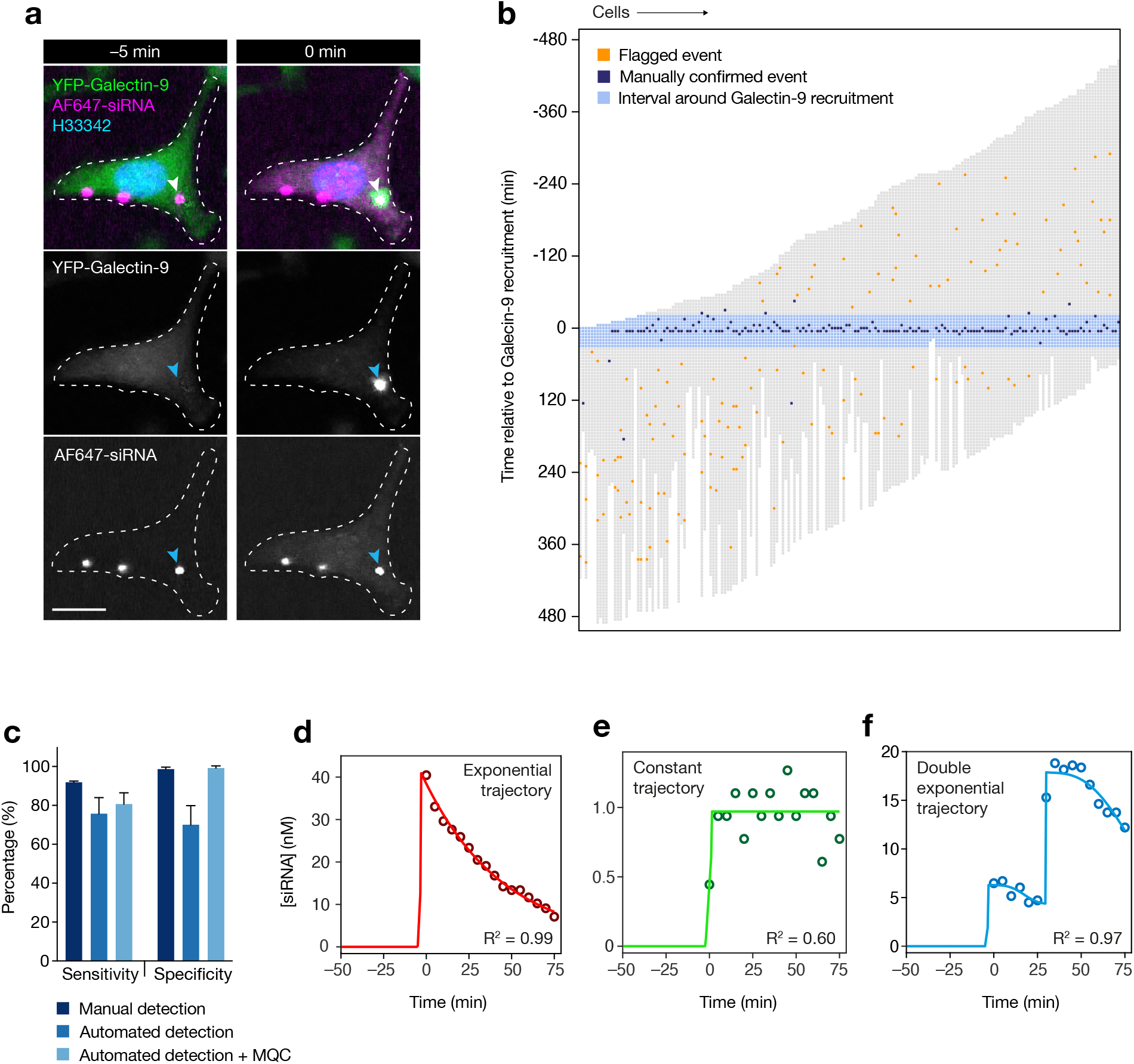
Automated detection and absolute quantification of cytosolic siRNA during lipid-mediated delivery. An Airyscan confocal detector was used to image HeLa cells expressing YFP-galectin-9 every 5 min during treatment with lipoplexed siRNA-AF647. **(a)** Image detail showing endosomal release and cytosolic dispersion of siRNA-AF647 in a single cell, coinciding with recruitment of YFP-galectin-9 to the lipoplex-containing vesicle (indicated by arrowheads). Images are representative of 187 release events from two independent experiments. Scalebar is 20 μm. **(b)** Trace-map of cells with recruitment of YFP-galectin-9 to vesicles containing siRNA-lipoplex, indicating the performance of the automated event detection in combination with manual quality control. Cell traces are organized in columns and aligned in time so that *t* = 0 is the first frame with detectable galectin-9 recruitment. *N* = 187 cells from 2 independent experiments. **(c)** Performance of manual and automated detection of endosomal siRNA release. Detection sensitivity and specificity was determined by comparing events indicated by galectin-9 recruitment and the manual or automated detection of siRNA-AF647 release to the cytosol. Manual quality control (MQC) was performed after automated event detection, to exclude false positive events. Mean ± s.d. is shown. *N* = 2 independent experiments. **(d-f)** Continuous monitoring of cytosolic siRNA-AF647 fluorescence intensity was translated into absolute concentration using condition-matched reference measurements. siRNA release magnitude estimations were made by fitting a mathematical model (lines) to the cytosolic siRNA concentration from single-cell measurements (circles). Examples are shown for the three different modelling approaches used, depending on the magnitude and kinetics of siRNA release: **(d)** a typical high-magnitude release event with exponential decay, **(e)** a high-noise low-magnitude release event (step-function), and **(f)**, two separate events occurring in quick succession.

We next sought to accurately quantify the cytosolic siRNA concentration after the release event. The typical cytosolic siRNA fluorescence intensity (median pixel) increases rapidly at the time of the release event and then gradually falls off, primarily as a consequence of cellular redistribution (Fig. 1f). The first frame with detectable cytosolic release most accurately corresponds to the released amount, because of the rapid, burst-like endosomal release and almost complete signal homogeneity within the cytosol. However, relying on a single frame for concentration measurements would result in noisy and potentially biased estimations. Therefore, we applied a mathematical model that captures the observed exponential decay of the cytosolic signal and fits the cytosolic signal intensities over 15 individual frames (75 min) to the model (Fig 2d–f, for details see Methods and Supplementary Note 1). This strategy enables unbiased estimations of absolute siRNA release amounts (expressed as a cytosolic concentration) for both low and high magnitude release events. We designed the model to allow for two release events to occur within 14 frames (70 min), and to consider this as a single release event with a magnitude of the sum of the two individual events (Fig. 2f). Additionally, the model fit (coefficient of determination, R^2^) provides a measure of accuracy and reliability of each individual cytosolic siRNA quantification. In summary, this acquisition and analysis strategy based on cytosolic siRNA fluorescence monitoring, enables sensitive, specific and quantitative siRNA release event detection.

As a model system to determine the intracellular dose-response of siRNA we selected a modified HeLa cell line stably expressing destabilized eGFP^18^ with a short half-life of ~48 min (Supplementary Fig. 5), providing a rapid read-out of knockdown effects from the eGFP fluorescence intensity. Conventionally, siRNA sequence potency can be measured by lipid transfection to determine the extracellular IC50. To evaluate if differences in extracellular IC50 between siRNA sequences is reflected in corresponding differences in the intracellular dose-response, we selected two siRNA sequences against eGFP with slightly different potency (relative IC50 siGFP-1: 0.29 nM, siGFP-2: 0.65 nM, absolute IC50 siGFP-1: 0.34 nM, siGFP-2: 0.92 nM), as measured with flow cytometry using defined extracellular siRNA concentrations (Fig. 3a). Similar IC50 for siGFP-1 was also obtained on mRNA level, when measured with RT-qPCR (Supplementary Fig. 6)

**Figure 3.**
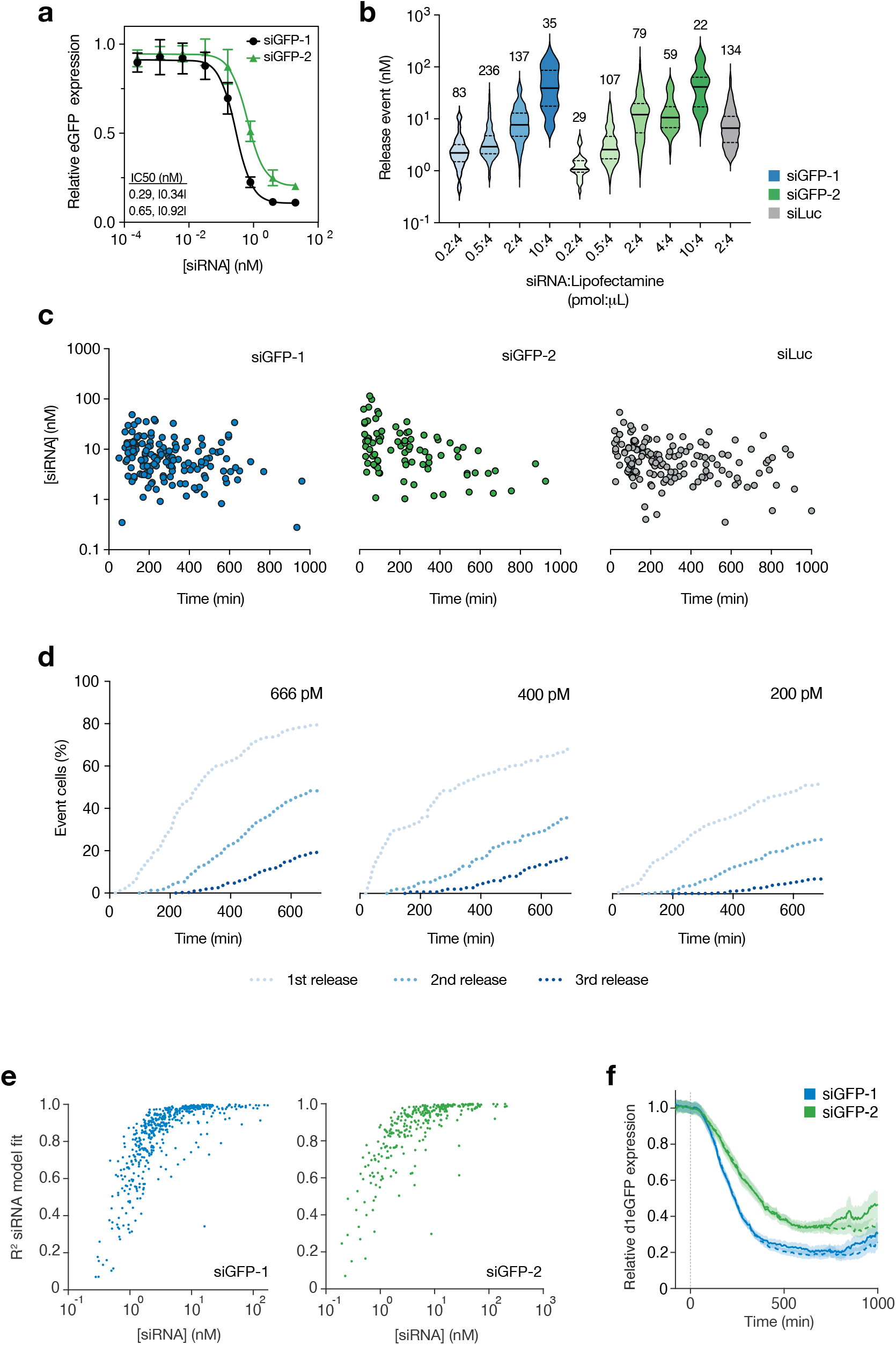
Magnitude of cytosolic delivery is determined by siRNA-to-lipid ratio. HeLa cells stably expressing d1-eGFP were treated with lipoplexed siRNA-AF647 targeting eGFP (siGFP-1 or siGFP-2) and analyzed by **(a)** flow cytometry or **(b-g)** live-cell microscopy followed by single-cell analysis**. (a)** d1-eGFP knockdown evaluated after 24 h siRNA-lipoplex incubation. Mean ± s.d. is shown. Relative and |absolute| IC50-values are indicated. *N* = 4 (siGFP-1) and 3 (siGFP-2) independent experiments. **(b)** Magnitudes of siRNA releases using lipoplexes formulated with different pmol siGFP-1 or siGFP-2 (constant for siLuc). Solid line is median and dashed lines are 25^th^ and 75^th^ percentiles. *N* = cells, indicated above violins. **(c,d)** Lipoplexes were formed with 2 pmol siRNA-AF647. For final concentrations, 16.7 μL, 10 μL or 5 μL prepared siRNA-lipoplex solution was used in experiments, to control the number of events. **(c)** Magnitude and time of siRNA release events during incubation with 200–666 pM lipoplexed siRNA-AF647. Start of experiment is *t* = 0. Single-cell measurements are shown, *N* = 151, 84, and 143 cells from three, one and four independent experiments, respectively. **(d)** Cumulative frequency of siRNA release events in cells incubated with indicated concentrations lipoplexed siGFP-1, siGFP-2 or siLuc. Mean values are shown. *N* = 2, 1 and 5 independent experiments, respectively. **(e,f**) Peak cytosolic siRNA concentrations estimated by release modelling and corresponding goodness-of-fit statistics (R^2^). Individual release events are shown. *N* = 546 and 316 events from 13 and 6 independent experiments, for siGFP-1 and siGFP-2, respectively. **(g)** d1-eGFP knockdown following one (solid lines) or multiple (dashed lines) siRNA release events per cell. Typically, after a second event is detected, subsequent measurements are excluded from analysis (solid lines). First event occurs at *t* = 0. Line is mean, shaded area is 80% confidence interval. *N* = 546 and 316 cells from 13 and 6 independent experiments for siGFP-1 and siGFP-2, respectively.

We then set out to determine experimental conditions that would provide a wide spectrum of different cytosolic siRNA release amounts, to use as the basis for subsequent dose-response analysis. The siRNA release amounts were similar for the two sequences as well as for an inactive control sequence (targeting luciferase, siLuc) but were highly dependent on the ratio of siRNA to transfection lipid (Fig. 3b). The release amounts of each individual release event, at a given ratio between lipid and siRNA, varied over more than an order of magnitude, possibly reflecting the heterogenous size of individual lipoplexes. The release magnitude was relatively independent of the time since addition of lipoplexes to the cells (Fig. 3c). Adding more lipoplexes to the cells (that is, adding a higher dose siRNA and a correspondingly higher dose transfection lipid at a constant ratio) resulted in more release events, occurring more rapidly (Fig. 3d) but with small effects on the average release magnitude (Supplementary Fig. 7). Importantly, very frequent events resulted in shorter cell-traces (in time) from the first to the second event. To obtain the intracellular dose-response relationship from a single intracellular siRNA dose, masking or disregarding expression data after the second event is necessary as this data would otherwise confound the dose-response determination. Thus, experimental conditions have to be optimized to achieve long analyzable traces before confounding second release events.

Based on the observations above, we collated multiple experiments with varying lipoplex doses and siRNA-to-lipid ratios for the two siGFP sequences incubated with HeLa-d1eGFP cells. We obtained a wide variety of release magnitudes with the quantification model fit (R^2^) typically above 0.75 for release magnitudes of 1 nM or more (Fig. 3e). Cellular eGFP intensities of tracked cells were corrected for bleaching, mitosis induced fluctuations and experiment-specific fluctuations as described in Methods (Supplementary Fig. 8–10). When averaging the eGFP expression of each individual cell experiencing a release event in this collection of data (with the time of the first release event set to 0 for each cell), knockdown was more prominent with the more potent siGFP-1 sequence compared to siGFP-2 (Fig. 3f). In addition, when comparing the eGFP expression of cells with or without masking of data after a second event, a longer knockdown duration can be appreciated in the non-masked data despite the experiments being optimized for few secondary release events. These secondary events happen at later timepoints during the monitoring, limiting the contribution to knockdown to primarily the end of the observation. In summary, evaluating eGFP expression after endosomal escape of variable amounts of siRNA (up until either a second release event or the cell is lost in tracking) can provide quantitative measures of knockdown kinetics for at least up to 17–20 h. A schematic of the key components of the analysis pipeline is shown in Supplementary Fig. 11.

We then turned to evaluating the dose-dependency of the eGFP knockdown. For this analysis, the cells were grouped into five quintiles depending on the model-based magnitude of siRNA release for both siGFP sequences, excluding very low-confidence siRNA quantifications (R^2^<0.3). The average cytosolic siRNA fluorescence after the release were similar to the average of the model estimates of siRNA release magnitude within each quintile, but with the fluorescence intensity consistently lower at high release magnitudes (Supplementary Fig. 12a, b). Cells exhibiting two closely spaced release events were accurately quantified by the model while the maximal magnitude of fluorescence intensity underestimated the total release amount, highlighting the advantage of the model-based quantifications (see an example of this effect in Fig. 2f). For siGFP-2, eGFP knockdown varied between 38% and 84% at 10 h after release of 0.93 (median, interquartile range 0.68–1.24) nM and 37.2 (22.2–55.8) nM (Fig. 4a and Supplementary Fig. 12). For the more potent siGFP-1 sequence knockdown was 72% after 10 h at both 1.21 (1.07–1.29) nM and 2.05 (1.97–2.23) nM. However, the accuracy of the siRNA quantifications was substantially worse in the lowest quintile (mean R^2^, quintile 1: 0.62, quintile 2: 0.80). Restricting analysis to only highly reliable quantifications (R^2^>0.75) yielded a clear dose-response relationship for both sequences with respect to knockdown induction kinetics, knockdown depth (nadir) and knockdown duration (Fig 4b). Thus, our live-cell monitoring of siRNA delivery and analysis pipeline can be used to elucidate the dose-response relationship of potent siRNA sequences.

We next wanted to determine the intracellular IC50 values for both siRNA sequences. As the degree of knockdown is dependent on time since release (Supplementary Movie 2), absolute intracellular IC50 values (that is, the concentration that results in 50% knockdown) will be highly dependent on the timepoint chosen. Relative IC50 values (the concentration with half-maximal inhibition) are potentially more stable. Plotting the eGFP expression level of single cells relative to the siRNA release amounts at various timepoints revealed comparatively stable relative IC50 values varying between 0.60–0.37 nM for siGFP-1 and 3.02–1.60 nM for the less potent siGFP-2 at 6 h and 10 h, respectively (Fig. 4c–e). Thus, the relative intracellular IC50 of an siRNA sequence is a measure of its potency and it correlates to conventionally measured extracellular IC50.

We finally wanted to obtain an intuitively understandable intracellular siRNA concentration or number of molecules that results in 50% knockdown for the two sequences respectively, that is, absolute IC50 values at the time point of maximal knockdown. For this, we observed that 10 h after release, knockdown was close to nadir or at a plateau for all release magnitudes for both sequences (Fig. 4b). This was also evident in the pooled data with release events of all magnitudes (Fig. 3g). At 10 h after release, 50% knockdown (absolute IC50) was determined to be 0.38 nM and 2.23 nM respectively for siGFP-1 and siGFP-2. However, relying solely on expression measurements at one time-point makes the estimation potentially overly dependent on the specific measurements at this time-point. To gauge the robustness the IC50 determinations we therefore adjusted a mathematical model to the dose-dependent knockdown kinetics, taking advantage of expression measurements from all time-points and all release magnitudes (Fig. 4f, see Methods). Based on this model, absolute IC50 at 10 h was estimated to 0.36 (0.14–0.65) nM for siGFP-1 and 3.36 (2.15–4.87) nM for siGFP-2 (median and 95% CI, calculated by bootstrapping), analogous to the values determined solely at 10h. Thus, we judge the absolute IC50 values determined at 10 h to be robust estimates for absolute intracellular IC50 values. Further, given the typical cytosolic volume of HeLa cells used in this study of 5000 fL (interquartile range 3800–5900 fL, Supplementary Fig. 13), we can estimate that 1100 siGFP-1 molecules are required for 50% knockdown at nadir (900 and 1400 molecules in cells with a size at the 25^th^ and 75^th^ percentile, respectively). For siGFP-2, 6700 molecules (5100 and 8000 molecules, in 25^th^ and 75^th^ percentile cells, respectively) are required for 50% knockdown.

**Figure 4.**
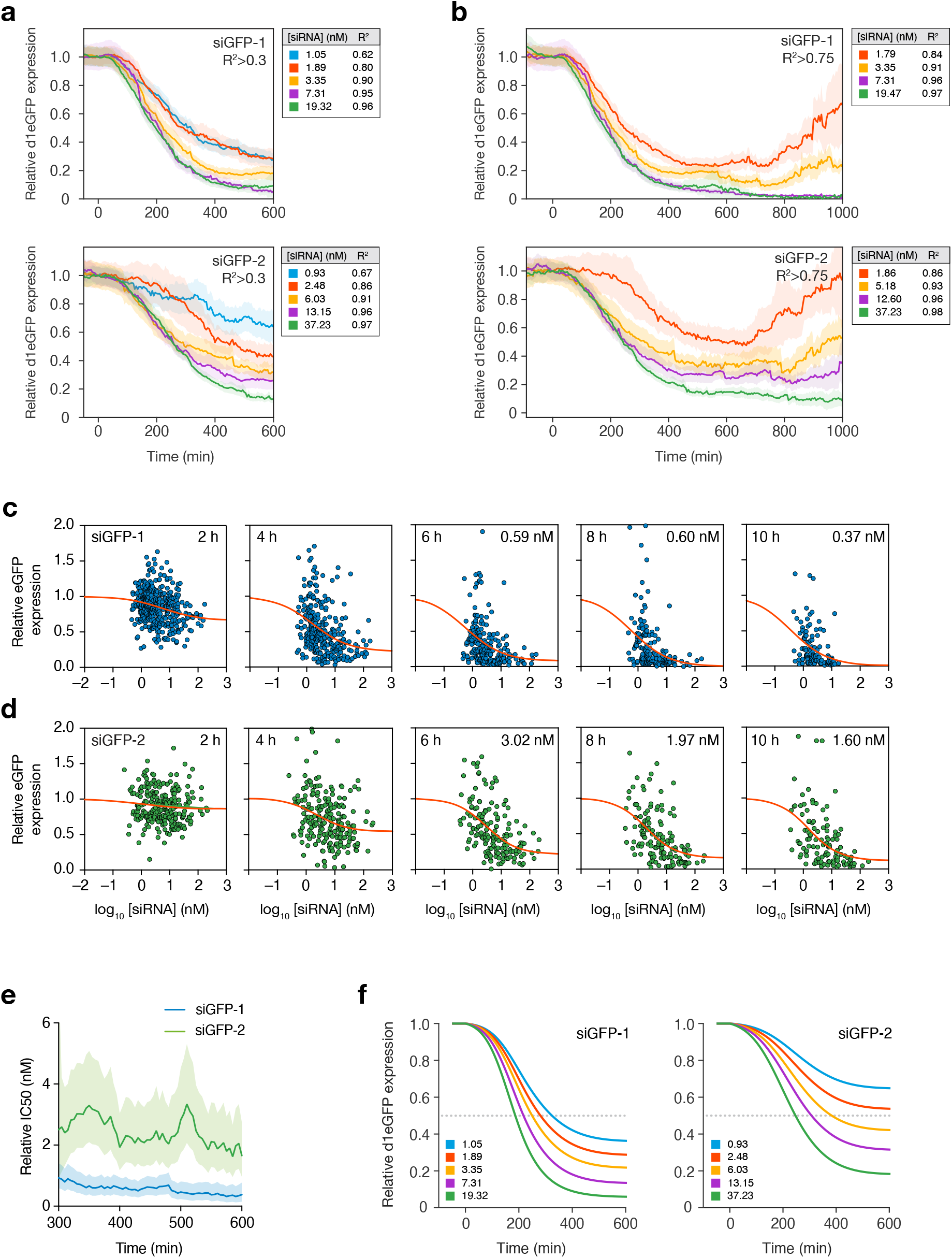
Single-cell knockdown kinetics are dose-dependent with respect to cytosolic siRNA amounts. HeLa cells stably expressing d1-eGFP were treated with 40–2000 pM lipoplexed siRNA-AF647 targeting eGFP (fixed siRNA-lipid ratio). A confocal microscope with Airyscan detector was used for live-cell imaging, followed by single-cell analysis. **(a,b)** Cells were ordered and divided in equal groups based on the model-estimated magnitude of siRNA release events. Traces are aligned so that *t* = 0 is the time of cytosolic siRNA detection, and d1-eGFP expression is shown relative to this time-point. Lines are mean eGFP expression per quantile, shaded areas are 80% confidence intervals. Median cytosolic siRNA concentration and mean R^2^-value per quantile is indicated. Model-fit (R^2^) thresholds of >0.3 **(a)** and >0.75 **(b)** were used. *N* ≥ 92, 58 (R^2^>0.3) 90 and 59 (R^2^>0.75) cells per quantile for siGFP-1 and siGFP-2, respectively. **(c,d)** Single-cell knockdown kinetics between 2 h and 10 h after siRNA release. Model-estimated peak cytosolic siRNA concentration and eGFP expression relative to *t* = 0 is shown per cell, for **(c)** siGFP-1 and **(d)** siGFP-2. Release events with model R^2^>0.3 are shown. Black line is interpolated sigmoidal curve fitted by least squares regression. Absolute knockdown IC50 (nM) is indicated. *N* between 2 h and (10 h) is ≤90 (≥50) and ≤50 (≥30) cells, for siGFP-1 and siGFP-2, respectively. **(e)** Relative knockdown IC50 for each time-point based on data shown in (c,d). Time from detected release event is indicated. Cells with release events with model R^2^>0.3 are shown. Line is IC50 and shaded area is 80% confidence intervals. **(f)** A model was used to estimate eGFP knockdown mediated by siGFP-1 or siGFP-2, predicted from the median peak cytosolic siRNA concentration after release in each quantile (indicated in graphs). All data shown is from 13 and 6 independent experiments, for siGFP-1 and siGFP-2, respectively.

## Discussion

Here we present a robust method to measure absolute cytosolic siRNA delivery amounts during lipid-mediated delivery. This method enabled us to determine single-cell intracellular dose-response relationships for siRNA knockdown of a reporter gene, including IC50, knockdown nadir and duration. We show that half-maximal knockdown induction and knockdown nadir is reached at a few hundred picomolar (~1000 intracellular molecules) with a potent siRNA sequence (siGFP-1). We also show that higher siRNA doses, beyond initial knockdown saturation prolong knockdown. The duration differ between cells having received ~13 and ~37 nM (~50 000 or ~100 000 molecules) of a less potent sequence (siGFP-2), highlighting a very large dynamic range in knockdown responses.

Previous cytosolic siRNA dose-response estimates have provided data at single time-points without information on cell-to-cell variability^19^. Cytosolic IC50 was estimated to ~2000-4000 molecules using electron microscopy of gold-labelled siRNA^9^ and through Ago2-immuno precipitation, IC50 was determined to be 10-110 RISC-loaded siRNA molecules (but with unclear RISC-loading efficiencies)^10^. As a comparison, using less accurate cytosolic concentration measurements (compared to this study), ~1.6 nM of siGFP-1 was determined to result in close to maximal knockdown^8^, similar to the almost maximal knockdown induction seen at above 3 nM here. While our initial aim was to establish the dose-response and in particular to measure the intracellular IC50, our knockdown kinetics analysis revealed that this is a dynamic concept – the IC50 values are strongly dependent on time since siRNA delivery (in addition to sequence potency). The nadir in eGFP expression is at 10–15 h after release for lower doses but this timing is dose-dependent and for very high siRNA releases, nadir does not seem to be reached at 20 h. The mechanism for this slow induction of full knockdown at high doses is not clear, but could involve a gradual increase in siGFP-associated RISCs when existing miRNA occupied RISCs continuously are replaced with newly synthesized RISCs. “Free” siRNA molecules, could thus act as a cytoplasmic depot of siRNA^11^, distinct from a recently demonstrated long-term non-cytoplasmic depot during GalNAc-siRNA delivery^20^.

An important aspect of our detection method is the ability to measure the reliability for each individual cytosolic siRNA quantification through the coefficient of determination, R^2^, of the fitted model. The reliability of the quantifications is thus demonstrated by both a high degree of model fit at estimated release amounts above 1 nM, and ultimately by the clear dose-response in knockdown for release events with high R^2^.

The Airyscan detector used here, is optimized for imaging within a small area in the center of the FOV of the confocal system, and only within this area the alignment is optimal for super-resolution imaging. In our experiments, we have used the full FOV of the microscopy system but without extracting any super-resolution information, essentially using the detector as a single, high dynamic range confocal detector with a 2.0 Airy unit pinhole. Other detection strategies are conceivable including modern large, high quantum-yield sCMOS-based, spinning-disk confocal systems with sparse pinhole patterns and high out-of-focus light suppression, but exploratory experiments indicated a higher detection limit for such a system. Various implementations of light sheet microscopy, for example, lattice light sheet^21^, HILO^22^, or other single-objective light sheet designs^23^ would be attractive from a detection limit perspective but would generally restrict FOV and experimental throughput.

In the present study the siRNA cargo is delivered with a cationic transfection lipid, resulting in larger discrete delivery events than what is achieved with clinical grade LNPs or siRNA-ligand conjugates. Detecting dispersed siRNA in the cytosol after a single LNP-delivery event would conceivably require a single-molecule detection strategy, which would be highly challenging given the substantial fluorescence from non-released particles. How multiple small release events occurring over time (such as from LNPs), in aggregate contribute to knockdown kinetics and potency is currently unclear, but the modelling strategies presented here offers a potential avenue to address this question.

Our method measures and estimates an apparent cytosolic concentration. However, this is a concentration over the space containing both the cytosolic fluid volume and all vesicles and other membrane enclosed organelles within the cytosolic space. Given that siRNA disperses homogenously initially after the release event, we assume the siRNA molecules are primarily free molecules within the cytoplasm. The effective siRNA concentration in this smaller volume (that is, only the fluid volume) is higher than the estimated concentration over the whole cytosolic space. The estimated number of released molecules is unaffected by this distinction.

This study is limited to determining the dose-response relationship of two siRNA sequences and a destabilized eGFP reporter gene in a single cell line. Future studies will elaborate how the intracellular dose-response varies between different cells, tissues, target genes and siRNA sequences. Indeed, recently it was suggested there are cell type specific variability in mRNA susceptibility and siRNA efficiency^24^, and mitotic activity^25^, target mRNA abundance^26^ and Ago2-expression^27^ is known to affect RNAi efficiency. The methodology presented here can be adapted for studies of other fluorescent reporter genes, fluorescent knock-in genes, and by using end-point analysis strategies (for example, fixation and immuno-staining) potentially any gene of interest.

A key challenge in efforts to understand and improve delivery methods for siRNA has been the unclear dose-response relationship of cytosolic siRNA, that we address here. Still unknown are the dose-responses for other nucleic acid therapeutics including mRNA and CRISPR compounds, of which the latter have the added complexity of being a multi-component nucleic acid mixture commonly consisting of a Cas9-mRNA, guide-RNA and a template-DNA all of which have distinct dose-response curves. Extension of the method presented here to these and other nucleic acid compounds has the potential to define dose-response relationships and facilitate rational efforts to improve the efficiency of this emerging class therapeutic molecules.

## Methods

### Cell culture and reagents

HeLa cells were ordered from American Type Culture Collection and verified to be free from mycoplasma contamination. Cells were cultured in Dulbecco’s Modified Eagle Medium (DMEM) (Hyclone, South Logan, UT, USA) supplemented with 10 % fetal bovine serum (FBS, Gibco), 2 mM glutamine (Thermo Fisher Scientific, Waltham, MA, USA), 100 U mL^−1^ penicillin, and 100 mg mL^−1^ streptomycin and incubated at 37°C, 5% CO_2_. Prior plating, cells were stained with trypan blue (Gibco) and counted with a Countess Automated Cell Counter (Invitrogen, Carlsbad, CA, USA) to obtain cell viability and concentration.

For all live-cell imaging experiments, cells were seeded 4–5 × 10^4^ cells per well in 8-well Lab-Tek II chambered cover glass slides (Nunc, Rochester, NY, USA) and incubated overnight. Before image acquisition, cells were washed with PBS, added with imaging medium (FluoroBrite DMEM (Gibco), 10% FBS, 2 mM glutamine, 2 mM HEPES) supplemented with 3.75 × 10^−3^ μg mL^−1^ Hoechst 33342 nuclear stain (Thermo Fischer Scientific) and further incubated for 1–2 hours.

Cycloheximide (CHX) and dimethyl sulfoxide (DMSO) were both from Sigma. Custom synthesized siRNA sequences were ordered from Integrated DNA Technologies. The following siRNA sequences were used: siGFP-1 sense: 5’-GGC UAC GUC CAG GAG CGC Atst-AF647-3’, siGFP-1 antisense: 5’-UGC GCU CCU GGA CGU AGC Ctst-3’, siGFP-2 sense: 5’-UGC UGC CCG ACA ACC ACU ACsC-AF647-3’, siGFP-2 antisense: 5’-UAG UGG UUG UCG GGC AGC AGsC-3’, siLuc sense: 5’-CUU ACG CUG AGU ACU UCG Atst-3’, siLuc antisense: 5’-CUU ACG CUG AGU ACU UCG Atst-AF647-3’. Lowercase denotes deoxynucleotides and ‘s’ indicate phosphorothioate linkage. Silencer Negative Control siRNA #1 (Invitrogen) was used as negative control for Real Time qPCR. Primer pairs for PCR were from Sigma (eGFP) and Invitrogen (GAPDH). The following primers were used: eGFP forward: 5’-ACG TAA ACG GCC ACA AGT TC-3’, eGFP reverse: 5’-AAG TCG TGC TGC TTC ATG TG-3, GAPDH forward: 5’-CTG GGC TAC ACT GAG CAC C-3’, GAPDH reverse: 5’-AAG TGG TCG TTG AGG GCA ATG-3’.

HeLa cells stably expressing d1-eGFP were established by transfecting cells with plasmid encoding d1-eGFP using Lipofectamine 2000 (Invitrogen) according to the manufacturer’s protocol. For selection, transfected cells were grown and sub-cultured in cell culture medium supplemented with 400 μg mL^−1^ G418 (Sigma), followed by single-cell fluorescence activated cell sorting using a BD FACSAria III cell sorter (Becton Dickinson, Franklin Lakes, NJ, USA) to obtain monoclonal cell lines. The d1-eGFP plasmid was from Invitrogen, and constructed by cloning the d1-eGFP synthetic gene into a pcDNA3.3-TOPO vector backbone.

### Lipoplex formulation

Formation of siRNA lipoplexes was performed using a fixed volume of Lipofectamine 2000 (LF2000) and variable siRNA concentrations, with a final siRNA-lipoplex solution volume of 100 μL. siRNA and LF2000 were first diluted in OptiMEM before mixing and incubating for 20 min at room temperature. The following pmol:μL ratio of siRNA to LF2000 was used for siRNA lipoplex formulation for microscopy experiments: siGFP-1: 0.2:4, 0.5:4, 2:4, 10:4; siGFP-2: 0.2:4, 0.5:4, 2:4, 4:4,10:4; siLuc: 2:4. For flow cytometry and RT-qPCR experiments, the following ratios were used: 2.6×10^−4^:4, 1.3×10^−3^:4, 6.4×10^−3^:4, 3.2×10^−2^:4, 0.16×10^−1^:4, 8×10^−1^:4, 4:4 and 20:4 pmol:μL siRNA to LF2000. For flow cytometry and RT-qPCR experiments, the added volume siRNA-lipoplex solution corresponded to 10% of the final volume in the well. For microscopy experiments, 5 μL, 10 μL, 16.7 μL or 50 μL of the prepared siRNA-lipoplex solution was added to microscopy slide wells, to control the number of lipoplexes internalized by cells. Since lipoplexes are formed before mixing with cell culture medium, adding varying volumes of the prepared siRNA-lipoplex solution will not change the particle size (that is, the number of siRNA-molecules in a single lipoplex), and hence the magnitude of siRNA release events will be equivalent (Supplementary Fig. 7). However, the final siRNA concentration (the number of lipoplexes) will differ depending on the volume of siRNA-lipoplex solution added.

### Live-cell imaging of siRNA release

HeLa cells stably expressing d1-eGFP or YFP-galectin-9 were plated in microscopy slides and incubated with imaging medium supplemented with 3.75×10^−3^ μg mL^−1^ Hoechst 33342. Cells were transferred to a preheated microscopy incubation chamber, and 4–6 positions with sparse and evenly distributed cells were selected. Immediately before starting image acquisition, lipoplexes formulated with siGFP-1, siGFP-2 or siLuc were added dropwise to the medium, to slow the particle settling rate and increase the likelihood of cells having single siRNA release events. Typically, 5 μL or 10 μL of the siRNA-lipoplex solution was added to the well, to control the final number of lipoplexes. For d1-eGFP knockdown experiments, one well was left untreated as control and imaged using the same settings as treated cells. Two *z*-plane images with 4 μm spacing were acquired per position with 5 min interval. The first *z*-plane was set to match the lower third of cells, to capture sudden endosomal siRNA release events. The second *z*-plane enabled capturing bright lipoplexes above the first imaging plane, that could potentially contribute to signal artifacts due to out-of-focus light. AF647-siRNA fluorescence was detected with an Airyscan detector while Hoechst33342 and d1-eGFP fluorescence were detected with a T-PMT detector. Typically, images were acquired for 12– 32 h for knockdown experiments, and 8 h for galectin-9 recruitment experiments.

### Single-cell tracking and quantification

After time-lapse image acquisition, measurements of siRNA and eGFP fluorescence were performed in each cell. D1-eGFP and Hoechst33342 images were denoised using the PureDenoise plugin^28^ in Fiji, to improve segmentation. In each image frame, individual cells were then segmented, tracked, masked, and measured in CellProfiler using customized analysis pipelines, as described below. Fluorescence measurements were performed on raw images. Analyses were performed in the lower *z*-plane if not stated otherwise. In brief, segmented nuclei were used for cell tracking and identified objects were labelled with unique identification numbers to structure single-cell measurements. Since boundaries of cells become increasingly difficult to identify correctly as the d1-eGFP expression decreases during knockdown, masks of segmented nuclei were also used for d1-eGFP fluorescence quantification. Comparisons of the d1-eGFP fluorescence intensity measured in nuclei alone and entire cells showed near identical results (Supplementary Fig. 8). Cytosolic AF647-siRNA fluorescence was measured with two types of segmentation masks, generated using cell boundaries identified in the d1-eGFP channel. A primary mask was used to detect sudden signal shifts, that is, for release event identification. A secondary mask, where the nuclear region was excluded, was used to quantify the cytosolic siRNA fluorescence intensity. In both cases, bright lipoplexes were identified in both imaging planes and excluded from the masks. Background signal intensity was calculated as the median pixel in the image after masking all identified objects (nuclei, cells and lipoplexes). All measurements were median pixel values calculated in the corresponding segmentation masks. Values were corrected for background intensity by subtraction. Data was exported from CellProfiler as Excel-files.

### Release event detection

A MATLAB script was used to automatically detect and validate sudden siRNA release events in single cells, using measurements from CellProfiler. In brief, the siRNA signal was evaluated in each frame to detect positive signal shifts. A shift in the signal intensity were considered as an siRNA release event if the shifted value and the five following measurements were all larger than the largest of i) the sum of the mean intensity value of the previous three frames and three times the standard deviation of the previous ten frames, or ii) the sum of the mean intensity value of the three previous frames and a fixed value. To reduce noise that might decrease the performance of the event-calling, a 5-frame moving median filter was applied to the siRNA signal before analysis. After automated detection, all identified events were validated manually. Regions-of-interest (ROIs) were created containing individual cells with detected events, that were then concatenated into image panels for inspection in Fiji. Events were classified as true or false. A detected event was considered true if siRNA release could be visually observed at the frame of detection or 10 frames ahead, and false if no visual release was observed. For true events, visual release typically coincided with automated detection of the event, or could be perceived 1–3 frames after automated detection. Cells showing signs of siRNA release before the first identified event (for example, redistribution of cytosolic siRNA, see 60-80 min in Fig. 1b) were excluded from subsequent analysis. If multiple release events occurred in the same cell, detected events were evaluated to the second true event.

### Sensitivity and specificity of cytosolic siRNA detection

HeLa cells stably expressing YFP-galectin-9 were imaged using live-cell microscopy for 8 h during treatment with lipoplex-formulated siGFP-1 at 2:4 pmol:μl ratio of siRNA to LF2000, adding 16.7 μL or 50 μL siRNA-lipoplex solution to the microscopy slide well. Endosomal siRNA release was then automatically and manually detected in each cell. For manual event detection, each cell was observed frame by frame until a release event was visible or until the end of the acquisition. For automated event detection, the quantification of cytosolic siRNA fluorescence intensity and signal shift detection was performed as described above. *De novo* recruitment and colocalization of galectin-9 with siRNA-lipoplexes was then manually evaluated in all cells, using the galecin-9 response as the true condition for evaluating release. Cells located partially outside the image border at the time of siRNA release or galectin-9 recruitment were excluded. Since evaluation of YFP-galectin-9 recruitment and localization is substantially more difficult following the first release event and initial galectin response, hampering the accuracy of the assay, only detection of the first galectin-9 verified events in each cell were analyzed. Likewise, with the manual and automated detection of cytosolic siRNA release, only the first detected event was evaluated. To determine the sensitivity and specificity of the cytosolic release detection to correctly identify the first siRNA release event in an evaluated cell, observations were classified as follows: Cytosolic release events detected within 5 frames before or after galectin-9 recruitment to the releasing lipoplex were considered true positive observations. If no cytosolic siRNA was detected even though galectin-9 was recruited to a visible lipoplex, the observation was considered false negative. Observations were classified as true negative if no cytosolic siRNA release was detected and no recruitment of galectin-9 to lipoplexes could be observed, and false positive if cytosolic release was detected in the absence of observable galectin-9 recruitment to the releasing lipoplex within 5 frames before or after the detection. A manual quality control was then performed of all siRNA release events identified by the automated detection algorithm, providing the opportunity to correct false positive events, that was then reclassified as true negative observations. This approach is analogous to the manual quality control of release events identified in experiments evaluating d1-eGFP knockdown (without galectin-9 reference), indicating similar detection sensitivity and specificity also in this setting. Release detection sensitivity was calculated as all true positive events divided by the sum of true positive and false negative observations. Specificity was calculated as true negative observations divided by the sum of true negative and false positive events.

### Cytosol volume calculation

HeLa cells expressing YFP-galectin-9 (used as a cytosolic marker) were used to calculate the average cytosol volume. Cells were prepared in microscope chambered glass slides, and nuclei stained with Hoechst 33342 as described above. Images were acquired using a confocal microscope, obtaining *z*-stacks with 12 *z*-plane spaced 1 μm apart containing the entire cell volume. Cells were manually outlined in Fiji to create ROIs, typically in the bottom, middle and upper *z*-plane of individual cells, and then using the “interpolate ROIs” function to interpolate ROIs in the remaining planes. This procedure was repeated for cell nuclei. For each experiment, 20 representative cells were analyzed. Cell area measurements (per *z*-plane) were exported as Excel-files. A MATLAB script was then used to calculate the volume of the cytosol by subtracting the volume of the nucleus from the total cell volume, given as femtoliters.

### Absolute quantification of intracellular siRNA

Before live-cell imaging experiments where intracellular siRNA was quantified, a calibration sample with 1 μM AF647-siRNA was prepared and imaged. Microscopy glass slides were prepared by adding 5 μL 1 μM AF647-siRNA diluted in cytosol-mimicking buffer (CMB) (15 g L^−1^ BSA, 125 mM KCl, 4 mM KH2PO4, 14 mM NaCl, 1 mM MgCl2, 20 mM HEPES), or 5 μL CMB only, to the center of slides. Droplets were covered with 18 × 18 mm 1.5# cover slips and imaged with the cover slip facing the objective. Three image stacks containing 30 *z*-planes with 0.5 μm interval spacing were acquired through the center of the sample using identical microscopy settings (for example, laser power, gain, pin-hole size, pixel size etc.) as used during live-cell imaging. A MATLAB script was then used to calculate the mean pixel intensity of images in the *z*-stack to determine the full width at half maximum (FWHM) interval of all *z*-planes (that is, the interval of *z*-planes with a mean pixel intensity equal or larger than the half maximum mean pixel intensity of a single *z*-plane image). A mean intensity projection across the *z*-dimension was created using all *z*-planes within the FWHM interval. A final mean reference image was then created from the z-projections of all three FWHM *z*-stacks. Pixel values were corrected for background fluorescence, calculated using the image stacks of CMB samples in the same way as for the siRNA samples. Cytosolic siRNA fluorescence, measured as the median pixel intensity in the cytosol and corrected for background fluorescence as described above, was corrected for uneven illumination (vignetting) and converted into absolute siRNA concentration by dividing measurements with the pixel value of the 1 μM siRNA reference (calibration) image at the center of the evaluated cell. The maximum siRNA value was used as a measure of the released siRNA quantity, except when model-based estimations were used (see below). The number of siRNA molecules per cell (*N*_C_) was calculated from the measured cytosol concentration (*C*) and measured typical cytosol volumes (*V*) as *N*_C_ = *C × V × N*_A_ where *N*_A_ is the Avogadro constant.

To evaluate the linearity of the fluorescence intensity readout of the microscopy setup, a reference curve was established using four-fold serial dilutions of AF647-siRNA between 1 nM and 1000 nM. The siRNA samples were diluted in CMB, prepared on microscopy glass slides, imaged in triplicates, and the fluorescence intensity was determined as described above.

### d1-eGFP half-life

HeLa-d1-eGFP cells were plated in microscopy slides and incubated with imaging medium supplemented with 3.75 × 10^−3^ μg mL^−1^ Hoechst 33342. Cells were transferred to a preheated microscopy incubation chamber and 4–6 positions with sparse and evenly distributed cells were located in two wells. Immediately before starting image acquisition, CHX or DMSO was added to either well with a final concentration of 50 μg mL^−1^ or 0.05%, respectively. For each position, 2 *z*-plane images were acquired every 5 min for 5 h. The d1-eGFP fluorescence intensity was quantified and analyzed as described below.

### Single-cell d1-eGFP expression analysis

After image acquisition, cells were tracked and quantified using CellProfiler as described above. A MATLAB script was then used for further analysis of single-cell measurements. In brief, cells with no or very low d1-eGFP expression in the beginning of the time-lapse acquisition were excluded from the analysis using a fixed value threshold. By evaluating changes in the size of cell nuclei, cells undergoing mitosis or apoptosis were identified. In cells where apoptosis was detected, measurements were masked starting 20 frames before cell death.

For d1-eGFP knockdown experiments, cells were excluded from analysis if they entered the field of view after the first 3 frames, to prevent including cells that may have had a prior undetected siRNA release event outside the frame. In addition, measurements after the detection of a second siRNA release event were masked, to limit all quantification of d1-eGFP knockdown to the effects of the first siRNA release event only. D1-eGFP measurements in cells with detected release events were corrected for bleaching and undetected release events, using measurements of cells in the same well that did not have any detected release event during lipoplex-treatment (Supplementary Fig. 9). Single-cell measurements were shifted in time so that all detected release events were aligned at *t* = 0 and corrected for mitosis-induced eGFP fluctuations using AF647-siLuc control experiments (for details, see Supplementary Fig 10. and Supplementary Note 2). Relative change in d1-eGFP expression was calculated by normalizing all values to the d1-eGFP intensity at the time of the siRNA release (*t* = 0). All cells were divided into equally sized quartiles or quintiles based on model-estimated magnitudes of siRNA release events.

For d1-eGFP half-life experiments with CHX, d1-eGFP signal was corrected for bleaching using measurements of DMSO-treated control cells. The relative change in d1-eGFP expression was calculated by normalizing single-cell measurements to the d1-eGFP intensity in the first acquired frame (*t* = 0).

### Modeling of siRNA release and knockdown kinetics

A full description and details of the mathematical model used to estimate siRNA release and d1-eGFP knockdown kinetics is available in Supplementary Note 2.

### Confocal microscopy

An inverted AxioObserver Z.1 LSM 710 epifluorescence confocal microscope with an Airyscan array detector unit (Carl Zeiss AG, Oberkochen, Germany) and equipped with a 40x Plan-Neofluar 1.3 numerical aperture (NA) oil-immersion objective was used for live-cell imaging acquisition. The pinhole was set to 2.15 Airy units (AU) in the software settings for the AF647-siRNA channel (effective pinhole used by the Airyscan detector is 1.25 AU), and fully opened (599 AU) for the d1-eGFP channel. The Airyscan Mode was Resolution vs. Sensitivity (R-S). A diode laser 405-30 (405 nm), a Lasos LGK 7812 argon laser (458 nm, 488 nm, 514 nm), and a HeNe633 laser (633 nm), were used as light source. A stage-top incubator with attached Temperature module, Heating Unit XL S and Heating Insert (Pecon), and CO_2_ control system was used for all live-cell imaging experiments, operating at 37 °C and 5% CO_2_. A Definite Focus module was used for auto-focus. The imaging system was operated under ZEN 2.1 (black edition).

### Flow cytometry

HeLa-d1-eGFP cells were seeded 3 × 10^4^ cells per well in a 48-well plate and incubated 24 h prior siRNA treatment with either siGFP-1, siGFP-2 or no treatment (control). Growth medium was removed and HEPES free imaging medium (FluoroBrite DMEM, 2 mM glutamine, 10 % FBS) was added to each well. A five-fold serial dilution of siRNA was performed in OptiMEM. Diluted siRNA was mixed with 4 μL LF2000 and incubated for 20 min at room temperature to form lipoplexes. A volume containing formed lipoplexes, corresponding to 10% of the final volume, was then added to the medium of each well, yielding final siRNA concentration between 0.26 pM and 20 nM with corresponding siRNA to LF2000 ratio between 2.6 × 10^−4^:4 and 20:4 pmol:μL. Cells were incubated for 24 h, washed with PBS, detached by trypsinization and resuspended in DMEM before transferring the medium to 12 × 75 polystyrene FACS tubes kept on ice. FACS tubes were centrifuged at 400 × *g* for 5 min, supernatant was decanted, and cells were resuspended in 0.5% BSA in PBS. The washing procedure was repeated, and cells resuspended in 0.5% BSA in PBS and analyzed with an Accuri C6 Flow Cytometer (Becton Dickinson, Franklin Lakes, NJ, USA). Cells were gated in side scatter/forward scatter plots and median fluorescence intensity was measured for each sample. HeLa wildtype cell fluorescence was subtracted to correct for background. Samples were performed in triplicates. Mean of triplicates were calculated and normalized to untreated HeLa-d1-eGFP cells.

### Real-Time qPCR

HeLa-d1-eGFP cells were seeded 1 × 10^5^ cells per well in a 12-well plate and incubated for 24 h prior siRNA treatment with siGFP-1 or negative control siRNA. Serial dilution of siRNA, lipoplex formulation and treatment was carried out as described above. Negative control siRNA was prepared at the same concentration as the highest siGFP dose. After 24 h lipoplex incubation, cells were washed with PBS and RNA was extracted using GenElute Mammalian Total RNA Miniprep Kit according to manufacturer’s protocol. Complementary DNA was obtained using SuperScript III First-Strand Synthesis System (Sigma) with random hexamer primers running on a MasterCycler EpGradient 5341 thermal cycler (Eppendorf AG, Hamburg, Germany). Real-Time qPCR was performed on a StepOnePlus Real-Time PCR System using MicroAmp Fast 0.1 mL 96-well Reaction Plates (Applied Biosystems, Foster City, CA, USA) and SYBR Green JumpStart Taq Readymix (Sigma) for the qPCR reactions. GAPDH was used as house-keeping gene, data was analyzed with StepOne Software v.2.3, and d1-eGFP expression fold-change relative to the control sample was calculated using the ΔΔC_t_ method.

### Software

CellProfileR^29^ (cellprofiler.org) version 2.2.0 was used to set up customized pipelines for fluorescence quantification in single cells. MATLAB 2016b was used for post-processing and data analysis. GraphPad Prism 9 version 9.0.1 was used for graphs and perform statistical testing. Fiji 2.1.0 and the plugin PureDenoise^28^ (https://bigwww.epfl.ch/algorithms/denoise) was used for image processing, data analysis, and image visualization. Illustrator 2020 23.0.2 was used for final figure design and composition.

### Statistics

Statistical tests and non-linear regressions were performed in GraphPad Prism 9. Non-linear regressions were fitted using “Sigmoidal, 4PL, X is log(concentration)” for sigmoidal curves, with Upper constraint = 1, Bottom constraint ≥0 with shared Hillslope when fitting multiple curves, and “Log-log line – X and Y both log” for standard curves.

## Data availability

All data supporting the findings of this study is available from the corresponding author upon reasonable request.

## Code availability

All custom computer code supporting the findings of this study is available from the corresponding author upon reasonable request.

## Acknowledgements

This work was supported by grants to A.W. from the Swedish Society for Medical Research (SSMF), the Gunnar Nilsson Cancer Foundation, the Mrs. Berta Kamprad Foundations, the Winklers Foundation, the Governmental funding of clinical research within the National Health Services (ALF), and the Wallenberg Center for Molecular Medicine, Lund University, and the Knut and Alice Wallenberg foundation.

## Author contributions

A.W. conceived the study and supported funding. H.H., H.D.R, and A.W. designed the experiments, analyzed the data and created figures. H.H. performed most of the experiments, H.D.R. and J.J. performed some of the experiments. H.D.R. designed software tools for image processing. L.H. assisted in siRNA sequence selection. J.W. designed mathematical models and contributed to data analysis. W.Z. provided technical assistance. H.H., H.D.R., J.W. and A.W. wrote the manuscript with input from all authors.

## References

1. Adams, D. et al. Patisiran, an RNAi Therapeutic, for Hereditary Transthyretin Amyloidosis. N. Engl. J. Med. 379, 11–21 (2018).

2. Balwani, M. et al. Phase 3 Trial of RNAi Therapeutic Givosiran for Acute Intermittent Porphyria. N. Engl. J. Med. 382, 2289–2301 (2020).

3. Garrelfs, S. et al. ILLUMINATE-A, A phase 3 study of lumasiran, an investigational RNAi therapeutic, in children and adults with primary hyperoxalurea type 1 (PH1). Nephrol. Dial. Transplant. 35, (2020).

4. Ray, K. K. et al. Inclisiran in Patients at High Cardiovascular Risk with Elevated LDL Cholesterol. N. Engl. J. Med. 376, 1430–1440 (2017).

5. Biscans, A. et al. Diverse lipid conjugates for functional extra-hepatic siRNA delivery in vivo. Nucleic Acids Res. 47, 1082–1096 (2019).

6. Hirsch, M. & Helm, M. Live cell imaging of duplex siRNA intracellular trafficking. Nucleic Acids Res. 43, 4650–4660 (2015).

7. Rehman, Z. Jr, Hoekstra, D. & Zuhorn, I. S. Mechanism of Polyplex- and Lipoplex-Mediated Delivery of Nucleic Acids: Real-Time Visualization of Transient Membrane Destabilization without Endosomal Lysis. ACS Nano 7, 3767–3777 (2013).

8. Wittrup, A. et al. Visualizing lipid-formulated siRNA release from endosomes and target gene knockdown. Nat. Biotechnol. 33, 870–876 (2015).

9. Gilleron, J. et al. Image-based analysis of lipid nanoparticle–mediated siRNA delivery, intracellular trafficking and endosomal escape. Nat. Biotechnol. 31, 638–646 (2013).

10. Stalder, L. et al. The rough endoplasmatic reticulum is a central nucleation site of siRNA-mediated RNA silencing. EMBO J. 32, 1115–1127 (2013).

11. Pei, Y. et al. Quantitative evaluation of siRNA delivery in vivo. RNA 16, 2553–2563 (2010).

12. Veldhoen, S., Laufer, S. D., Trampe, A. & Restle, T. Cellular delivery of small interfering RNA by a non-covalently attached cell-penetrating peptide: quantitative analysis of uptake and biological effect. Nucleic Acids Res. 34, 6561–6573 (2006).

13. Laufer, S. D., Recke, A. L., Veldhoen, S., Trampe, A. & Restle, T. Noncovalent Peptide-Mediated Delivery of Chemically Modified Steric Block Oligonucleotides Promotes Splice Correction: Quantitative Analysis of Uptake and Biological Effect. Oligonucleotides 19, 63–80 (2009).

14. Landesman, Y. et al. In vivo quantification of formulated and chemically modified small interfering RNA by heating-in-Triton quantitative reverse transcription polymerase chain reaction (HIT qRT-PCR). Silence 1, 16 (2010).

15. Huff, J. The Airyscan detector from ZEISS: confocal imaging with improved signal-to-noise ratio and super-resolution. Nat. Methods 12, i–ii (2015).

16. Thurston, T. L. M., Wandel, M. P., von Muhlinen, N., Foeglein, Á. & Randow, F. Galectin 8 targets damaged vesicles for autophagy to defend cells against bacterial invasion. Nature 482, 414–418 (2012).

17. Du Rietz, H., Hedlund, H., Wilhelmson, S., Nordenfelt, P. & Wittrup, A. Imaging small molecule-induced endosomal escape of siRNA. Nat. Commun. 11, 1–17 (2020).

18. Li, X. et al. Generation of Destabilized Green Fluorescent Protein as a Transcription Reporter *. J. Biol. Chem. 273, 34970–34975 (1998).

19. Yang, N. J. et al. Cytosolic delivery of siRNA by ultra-high affinity dsRNA binding proteins. Nucleic Acids Res. 45, 7602–7614 (2017).

20. Brown, C. R. et al. Investigating the pharmacodynamic durability of GalNAc–siRNA conjugates. Nucleic Acids Res. 48, 11827–11844 (2020).

21. Chen, B.-C. et al. Lattice light-sheet microscopy: Imaging molecules to embryos at high spatiotemporal resolution. Science 346, (2014).

22. Tokunaga, M., Imamoto, N. & Sakata-Sogawa, K. Highly inclined thin illumination enables clear single-molecule imaging in cells. Nat. Methods 5, 159–161 (2008).

23. Sapoznik, E. et al. A versatile oblique plane microscope for large-scale and high-resolution imaging of subcellular dynamics. eLife 9, e57681 (2020).

24. Ferguson, C. M., Echeverria, D., Hassler, M., Ly, S. & Khvorova, A. Cell Type Impacts Accessibility of mRNA to Silencing by RNA Interference. Mol. Ther. - Nucleic Acids 21, 384–393 (2020).

25. Bartlett, D. W. & Davis, M. E. Insights into the kinetics of siRNA-mediated gene silencing from live-cell and live-animal bioluminescent imaging. Nucleic Acids Res. 34, 322–333 (2006).

26. Hong, S. W., Jiang, Y., Kim, S., Li, C. J. & Lee, D. Target Gene Abundance Contributes to the Efficiency of siRNA-Mediated Gene Silencing. Nucleic Acid Ther. 24, 192–198 (2014).

27. Vickers, T. A., Lima, W. F., Nichols, J. G. & Crooke, S. T. Reduced levels of Ago2 expression result in increased siRNA competition in mammalian cells. Nucleic Acids Res. 35, 6598–6610 (2007).

28. Luisier, F., Vonesch, C., Blu, T. & Unser, M. Fast interscale wavelet denoising of Poisson-corrupted images. Signal Process. 90, 415–427 (2010).

29. Kamentsky, L. et al. Improved structure, function and compatibility for CellProfiler: modular high-throughput image analysis software. Bioinformatics 27, 1179–1180 (2011).

