## Supplementary information for "Absolute quantification and single-cell dose-response of cytosolic siRNA delivery"

Manuscript archived at bioRxiv April 21, 2021

**Supplementary Figures**

| | |
|---|---|
| Supplementary Fig. 1 | Linear relationship between AF647-siRNA fluorescence and concentration. |
| Supplementary Fig. 2 | Image vignetting at maximal FOV. |
| Supplementary Fig. 3 | Cells tracked through a series of time-lapse microscopy frames. |
| Supplementary Fig. 4 | Automated siRNA releases event detection. |
| Supplementary Fig. 5 | Half-life of d1-eGFP protein. |
| Supplementary Fig. 6 | Knockdown IC50 from d1-eGFP-mRNA expression. |
| Supplementary Fig. 7 | Release event magnitude is not affected by final siRNA dose. |
| Supplementary Fig. 8 | Nucleus and cell mask quantification. |
| Supplementary Fig. 9 | eGFP fluorescence baseline drift correction. |
| Supplementary Fig. 10 | Correction of mitosis induced fluctuations. |
| Supplementary Fig. 11 | Process and analysis pipeline for single cell RNAi kinetics. |
| Supplementary Fig. 12 | Absolute quantifications of siRNA release events in single cells. |
| Supplementary Fig. 13 | Cytosol volume of a typical HeLa cell. |
| Supplementary Fig. 14 | Trajectories used for fitting siRNA release model |

**Supplementary Notes**

| | |
|---|---|
| Supplementary Note 1 | Modeling cytosolic siRNA release and eGFP knockdown response |
| Supplementary Note 2 | Apparent eGFP expression increase during mitosis |

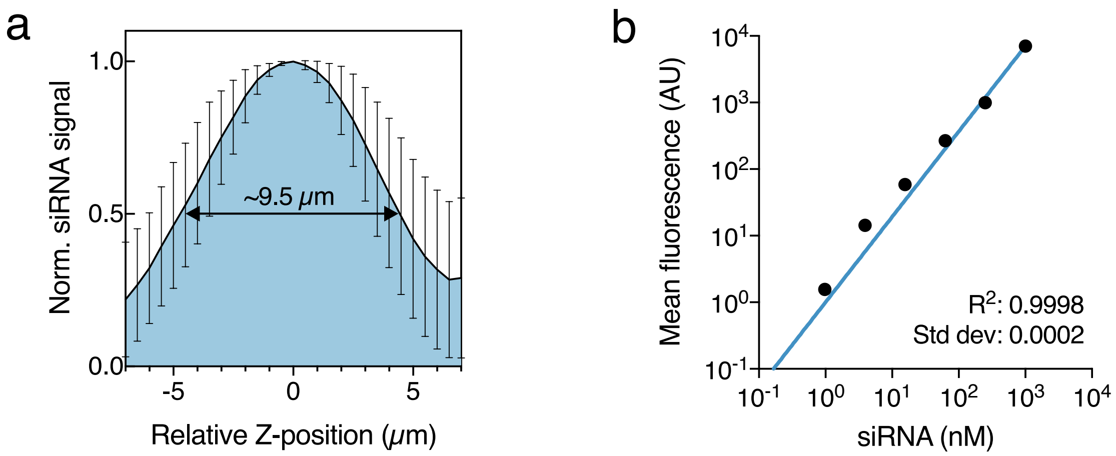


**Supplementary Fig. 1 | Linear relationship between AF647-siRNA fluorescence and concentration.** AF647-siRNA was prepared in cytosolic mimicking buffer on microscopy glass slides and imaged using an Airyscan confocal detector. (**a**) siRNA fluorescence intensity through a 1000 nM solution with a thickness similar to an average cell. Two-headed arrow indicate full-width at half maximum (FWHM). Mean ± s.d. from 21 independent experiments. (**b**) Mean siRNA fluorescence intensity within FWHM for 1–1000 nM serial dilutions. Log-log linearity between siRNA fluorescence and concentration were assessed per experiment. Representative data from one experiment is shown. Indicated R2-value and s.d. are means of three independent experiments.

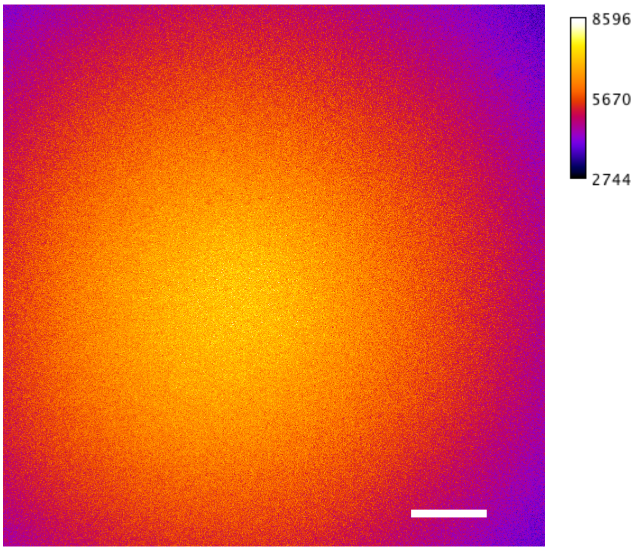


**Supplementary Fig. 2 | Image vignetting at maximal FOV.** Reference z-stacks of a 1000 nM AF647-siRNA reference slide were acquired with an Airyscan detector before live-cell imaging. Image vignetting at maximal field of view is illustrated. Scalebar is 50 μm. Data is representative of 21 independent experiments.

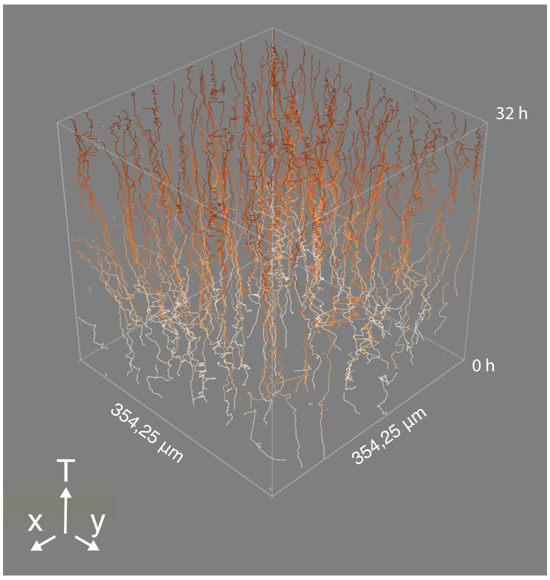


**Supplementary Fig. 3 | Cells tracked through a series of time-lapse microscopy frames.** Identified nuclei are connected between frames to facilitate tracking and measuring individual cells. Tracks are shown as lines proceeding along the T-dimension.

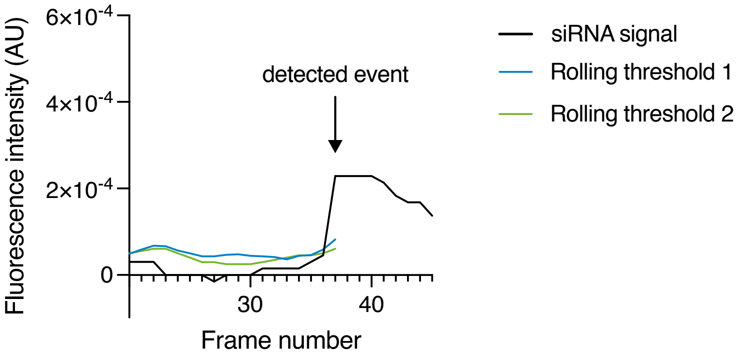


**Supplementary Fig. 4 | Automated siRNA release event detection.** An algorithm for automated detection of siRNA release was used, to identify cytosolic release events above one of two prespecified rolling thresholds (see Methods for details). Representative cytosolic siRNA measurements from one cell is shown, with a detected release event at frame 37.

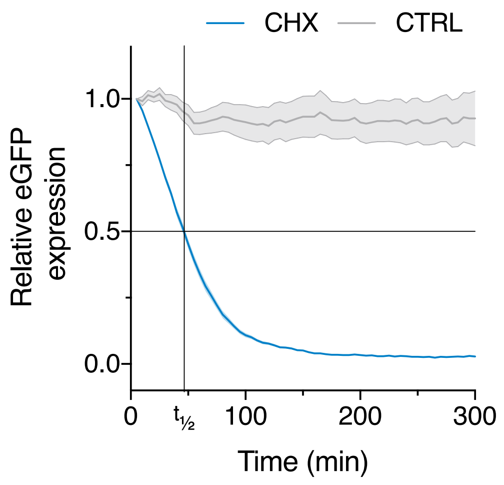


**Supplementary Fig. 5 | Half-life of d1-eGFP protein.** HeLa cells expressing d1-eGFP were imaged every 5 min during treatment with 50 μg mL–1 cycloheximide (CHX) or DMSO (control). The d1-eGFP expression is shown relative to t = 0 (start of acquisition immediately after addition of CHX or DMSO). D1-eGFP half-life (t1/2) was estimated to ~48 min. Mean and 95% confidence interval (shaded area) is shown. N = 218 and 102 cells for CHX and DMSO, respectively, from three independent experiments.

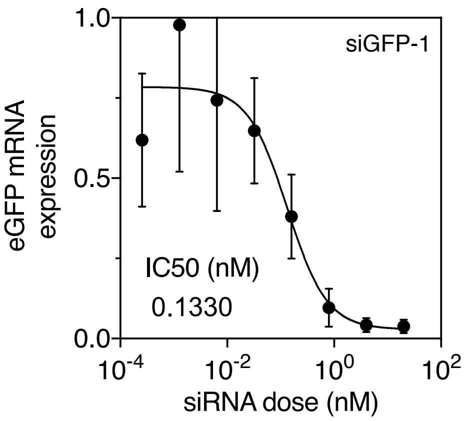


**Supplementary Fig. 6 | Knockdown IC50 from d1-eGFP-mRNA expression.** HeLa cells expressing d1-eGFP were treated with the indicated concentrations lipoplexed siGFP-1 for 24 h, and eGFP-mRNA expression was analyzed by RT-qPCR. Mean ± s.d. is shown. A sigmoidal dose-response curve was fitted with non-linear regression, to estimate relative knockdown IC50 (indicated). N = 4 independent experiments.

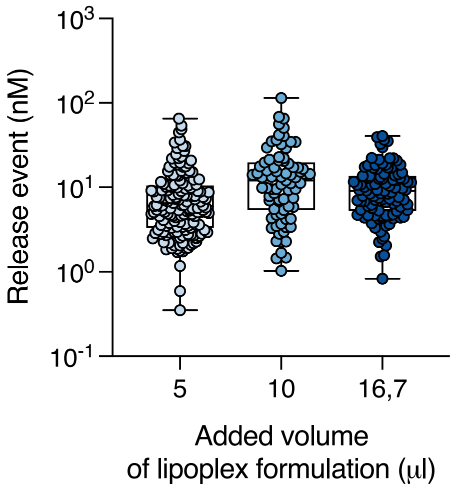


**Supplementary Fig. 7 | Release event magnitude is not affected by final siRNA dose.** HeLa cells expressing d1-eGFP were incubated with lipoplexed siRNA (siGFP-1, siGFP-2, or siLuc) formed with a fixed siRNA-lipid ratio (2 pmol:4 μL). To control the number of release events, the volume of prepared siRNA-lipoplexes added to cells were adjusted between 5–16.7 μL. A confocal microscope with Airyscan detector was used for live-cell imaging, followed by single-cell analysis. The siRNA release magnitude in individual cells is shown for each condition. Box-plot shows median with 25th and 75th percentiles, bars indicate min and max. N = 166, 79 and 105 cells from two, one, or five independent experiments, respectively.

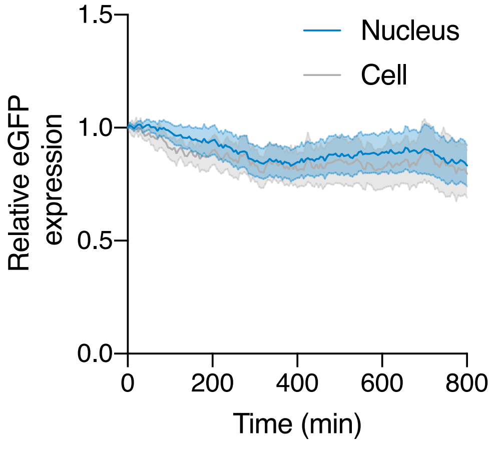


**Supplementary Fig. 8 | Nucleus and cell mask eGFP quantification.** HeLa cells expressing d1-eGFP were imaged every 5 min with a confocal microscope without treatment. D1-eGFP fluorescence intensity was measured using masks based on cell boundaries (entire cells) or cell nuclei only, and is shown relative to t = 0 (start of acquisition). Line is mean, shaded are is 95% confidence interval. N = 131 cells from three independent experiments.

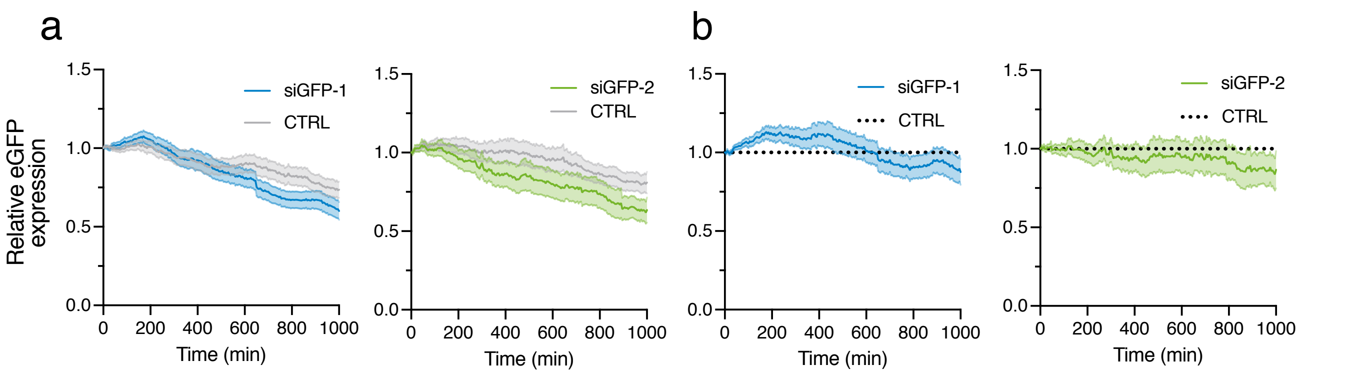


**Supplementary Fig. 9 | eGFP fluorescence baseline drift correction.** HeLa cells expressing d1-eGFP incubated with lipoplexed AF647-siRNA (siGFP-1 or siGFP-2, siRNA-to-lipid ratio 0.2–10 pmol:4 µL), or no treatment (CTRL), were imaged every 5 min with a confocal microscope. (**a**) eGFP expression was quantified in control cells and siRNA-treated cells with no detectable cytosolic release events, to evaluate knockdown from release events below the detection limit. (**b**) eGFP expression in siRNA-treated cells normalized to the mean of control cells, shown relative to t = 0 (start of acquisition). Line is mean, shaded area is 95% confidence interval. N = 362, 138 (siRNA), 427 and 267 (control) cells for siGFP-1 and siGFP-2, respectively, from 7 and 5 independent experiments. During standard post-acquisition processing, eGFP expression in siRNA-treated cells without detectable release events were routinely used to correct for bleaching and knockdown caused by non-detectable events.

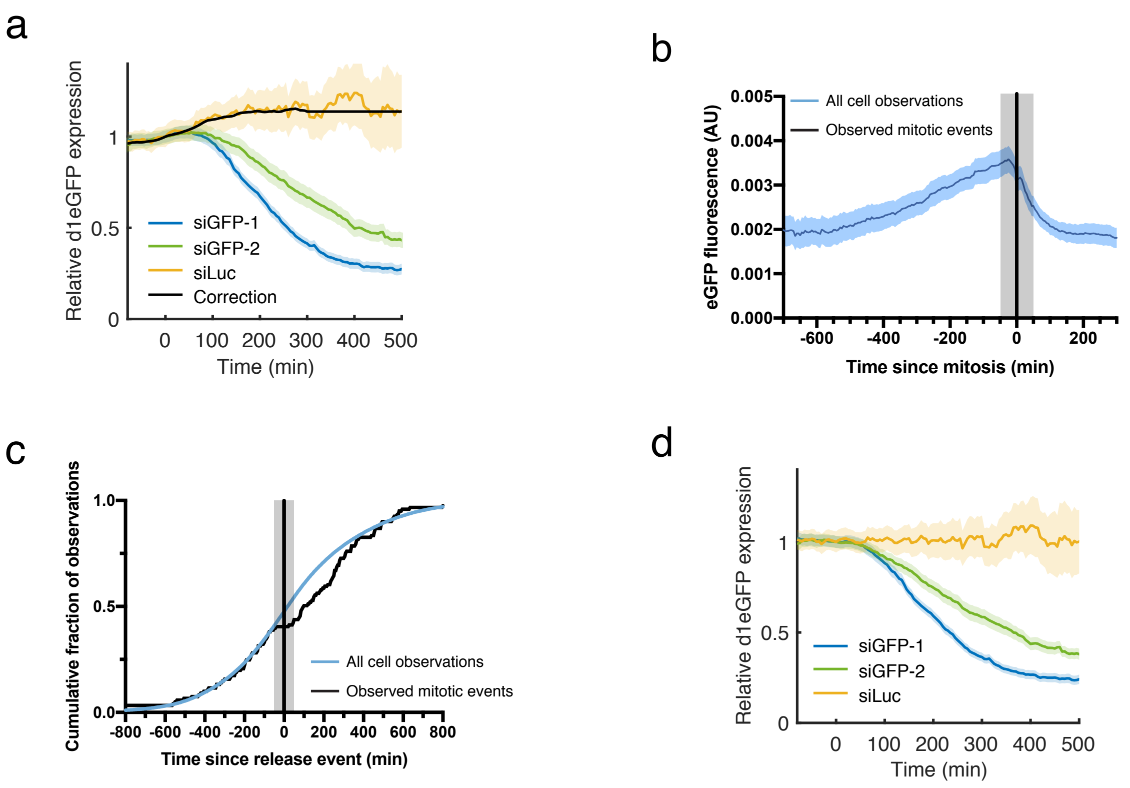


**Supplementary Fig. 10 | Correction of mitosis-induced eGFP fluctuations.** HeLa cells expressing d1-eGFP were treated with lipoplexed AF647-siRNA, acquiring images every 5 min with an Airyscan confocal detector followed by single-cell analysis. (**a**) eGFP expression after siRNA release events of either siGFP-1, siGFP-2 or inactive siLuc. Release event occurs at t = 0. Lines are mean, shaded areas are 80% confidence intervals. Black line is 10-frame moving average of siLuc. (**b**) eGFP expression during mitosis (t = 0) in cells with no siRNA release event. Shaded area indicates 5 frames before and after mitosis. N = 181 cells. (**c**) Cumulative distribution of observed mitotic events (black line) compared to the cumulative distribution of all cell observations (blue line), in cells exhibiting a release event at t = 0. N = 389 cells. (**d**) eGFP expression after correction using the moving average eGFP fluorescence in cells with siLuc release events. Data is based on 13, 6 and 4 independent experiments for siGFP-1, siGFP-2 and siLuc respectively.

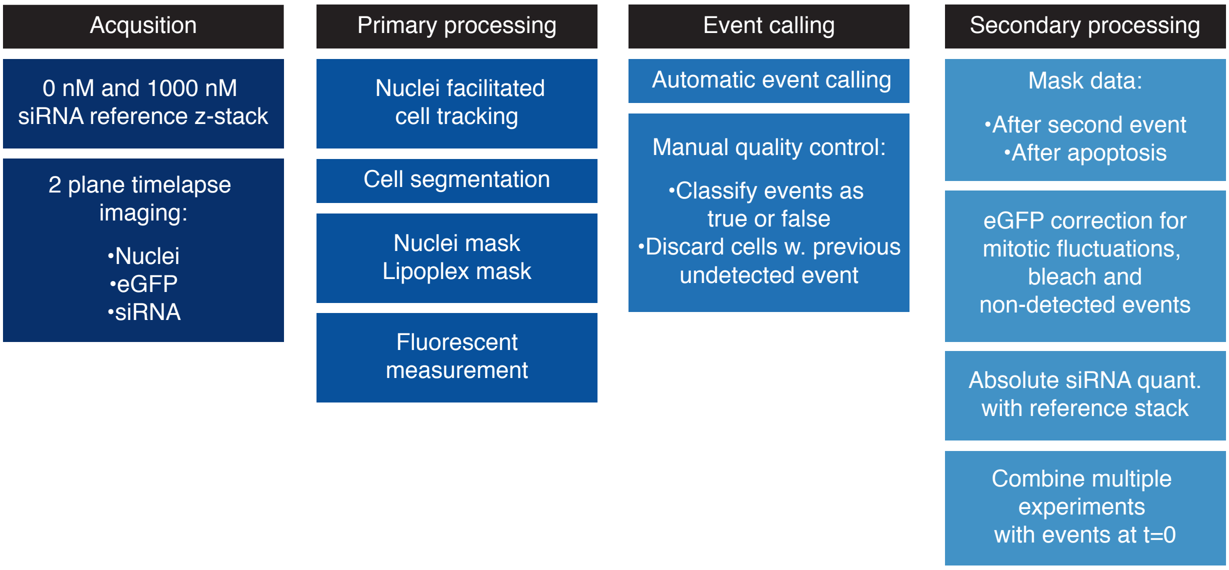


**Supplementary Figure 11 | Processing and analysis workflow for single-cell RNAi kinetics.** Flowchart showing the post-acquisition processing and analysis workflow for determining single-cell eGFP knockdown kinetics and absolute quantification of cytosolic siRNA concentration.

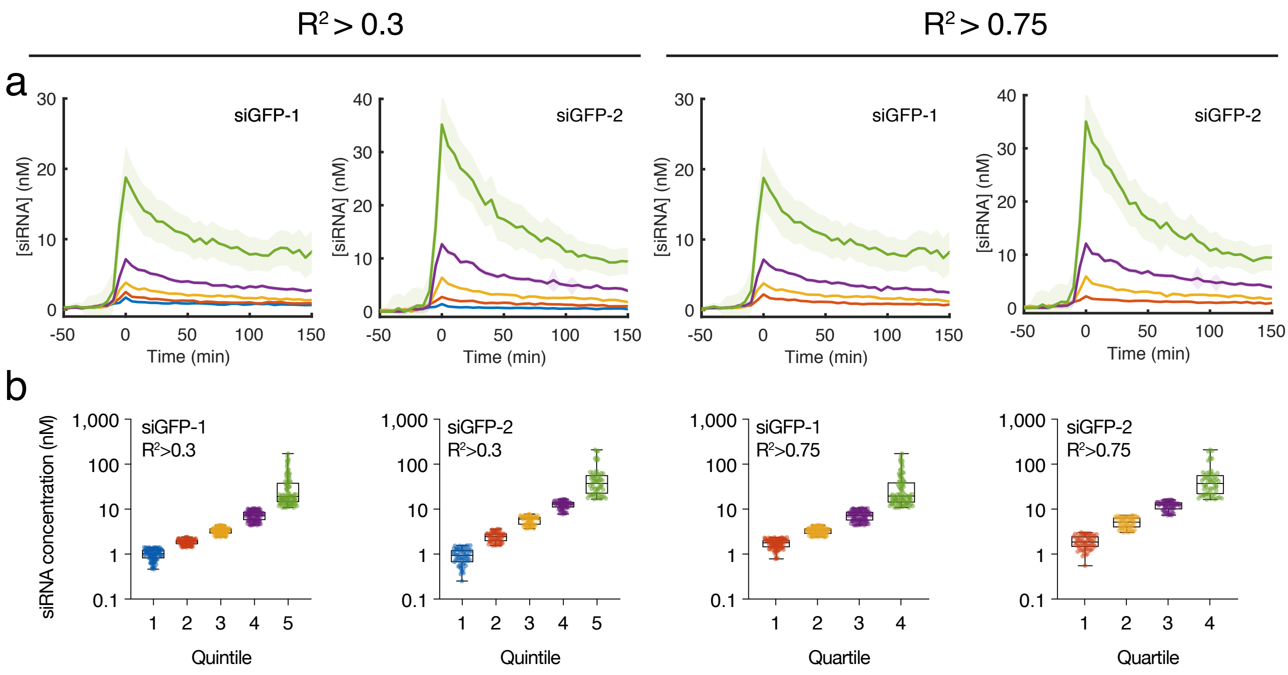


**Supplementary Fig. 12 | Absolute quantifications of siRNA release.** HeLa cells stably expressing d1-eGFP were treated with 40–2000 pM lipoplexed AF647-siRNA targeting eGFP (fixed siRNA-lipid ratio). A confocal microscope with Airyscan detector was used for live-cell imaging, followed by single-cell analysis. Cells were ordered and divided in equal groups based on the model-estimated magnitude of siRNA release events. (**a**) Lines are mean cytosolic siRNA concentration (measured) per quantile, shaded areas are 95% confidence intervals. Traces are aligned so that t = 0 is the time of cytosolic siRNA detection. Model-fit (R2) thresholds of >0.3 and >0.75 were used. (**b**) Model-estimated peak cytosolic siRNA concentration per cell is shown. Box-plots indicate median with 25th and 75th percentile, bars indicate min and max. N ≥ 92, 58 (R2>0.3) 90 and 59 (R2>0.75) cells for siGFP-1 and siGFP-2, respectively, from 13 and 6 independent experiments.

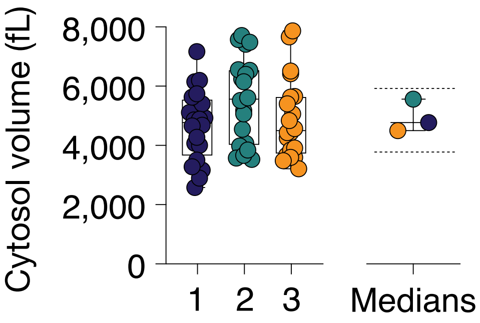


**Supplementary Fig. 13 | Cytosol volume of a typical HeLa cell.** HeLa cells expressing a cytosolic marker (YFP-galectin-9) were imaged using confocal microscopy, and the cytosol volume was calculated from measurements of the entire cell and nucleus volume. N = 60 cells from three independent experiments (numbered 1–3), and medians of cells per experiments are shown. Dashed lines indicate mean 25th and 75th percentiles.

### Supplementary Note 1 —Modeling cytosolic siRNA release and eGFP knockdown response

#### 1 siRNA model fitting

Here, we describe how the peak cytosolic siRNA fluorescence intensity is estimated by modeling release events. If observations were exact and continuous, the highest measured value of the siRNA quantity would have been the most accurate estimate of the peak siRNA concentration. However, since siRNA is observed with measurement error and only at discrete time-points, we cannot obtain this data directly from observations. Instead we build a mathematical model for the data, which in turn provides an estimated maximal value of the siRNA amount, which we denote $\widehat{siRNA}$,

The data on which the model is built is the following: For each cell, we have observations of the siRNA fluorescence intensity, at frames $-10$ to 15 relative to the release event occurring at frame 0.

Three different types of distinct trajectories of the change in siRNA fluorescence intensity are fitted to this data. The trajectories were selected based on the observed kinetics of siRNA release and redistribution. First is the constant trajectory, where a sudden signal increase is observed at the release, which then levels off to a constant (Fig. 2e). Second is the exponentially decaying trajectory (Fig. 2d). Third is the case with two subsequent release events (double exponential) (Fig. 2f).

When examining siRNA measurements manually, it is often possible to see which type of event has occurred. To have the model autonomously decide what trajectory is most appropriate, all three trajectories are fitted for each cell. The model with the best fit is then selected using Bayesian information criteria (BIC) [1]. The criteria will punish models using too many parameters, thus penalizing overfitting. This is required as the double trajectory will typically give a better fit than the single trajectory, making simpler measures for evaluating goodness-of-fit (like $R^2$) disadvantageous, since it would always select the most complex model.

##### 1.1 Functions

Here we mathematically define the three trajectories used. All three describe a fast linear growth after the first siRNA release, then followed by different behaviours. The trajectories are denoted $f(t;\boldsymbol{\theta})$ where the first argument is time and second argument is parameters.

###### 1.1.1 Constant trajectory

The function which we fit is the following, consisting of two terms

$$f_1(t;\boldsymbol{\theta}) = \mathbb{I}\left(t > \theta_1\right)\theta_2 + \mathbb{I}\left(\theta_3 < t < \theta_1\right)\theta_2 \frac{(t-\theta_3)}{\theta_1}.$$

The second term describes the linear growth from the release event (at frame $\theta_3$) until frame having the largest value, and the first term describes the trajectory after the growth is complete.

Here the three parameters represent $\theta_1-$ time after which the linear growth is complete, $\theta_2-$ the value after the linear growth is complete and $\theta_3-$ time where the linear growth starts. For this trajectory $\widehat{siRNA} = \theta_2$.

##### 1.1.2 Exponential trajectory

The function which we fit is the following function consisting of two terms.

$$f_1(t;\boldsymbol{\theta}) = \mathbb{I}\left(t > \theta_1\right)\left(\theta_2 + \exp\left(\theta_4 - \frac{(t-\theta_1)^{\theta_6}}{\theta_5}\right)\right) + \mathbb{I}\left(\theta_3 < t < \theta_1\right)\left(\theta_2 + \exp(\theta_4)\right)\frac{(t-\theta_3)}{\theta_1}.$$

Here we have the same parameters as the constant function, with the addition of three more: $\theta_4-$ logarithm of the largest value from which the exponential function decays. $\theta_5-$ scale on which the exponential function decays, $\theta_6-$ shape of the exponential function. For this function $\widehat{siRNA} = \theta_2 + \exp(\theta_4)$.

##### 1.1.3 Double exponential trajectory

This function is identical to the exponential trajectory except that it has one additional growth with exponential decay after the initial event, at a time point which needs to be estimated by the model. Thus, in total it has ten parameters.

#### 1.2 Fitting

For each cell, functions are fitted using least squares (LS) estimate:

$$\hat{\boldsymbol{\theta}} = \arg\min_{\boldsymbol{\theta}} \sum_{i=-10}^{15} \left(y_i - f_j(i;\boldsymbol{\theta})\right)^2,$$

for the three trajectories $f_1(t;\boldsymbol{\theta}), f_2(t;\boldsymbol{\theta})$ and $f_3(t;\boldsymbol{\theta})$. As these functions are non convex (i.e. with potentially multiple local maxima), ten different random starting points for $\boldsymbol{\theta}$ is used for each function, and the parameter with the smallest LS value is selected. Examples of the different types of functions found are illustrated in Supplementary Fig. 14. After finding the function with the best obtained fit for each trajectory type, we use BIC to select the best of the three.

#### 1.3 Model accuracy

For the best model, here denoted $f$, we also calculate what proportion of the variance that is explained by the fitted function, i.e. $R^2$. This statistic is calculated as follows

$$\tilde{R}^2 = 1 - \frac{\sum_{i=-10}^{15}\left(y_i - f(i;\hat{\boldsymbol{\theta}})\right)^2}{\sum_{i=-10}^{15}\left(y_i - \bar{y}\right)^2},$$

where $\bar{y} = \frac{1}{25}\sum_{i=-10}^{15} y_i$.

#### 1.4 Example

In Supplementary Fig. 14, the three different trajectories and all fits found for one release event is shown. In this instance, the exponential trajectory had the lowest BIC and hence was chosen. Notice that the double exponential trajectory had the best fit ($R^2$). However, from examination of the data there is no clear evidence of two release events. This is correctly identified by the BIC and the simpler trajectory is selected.

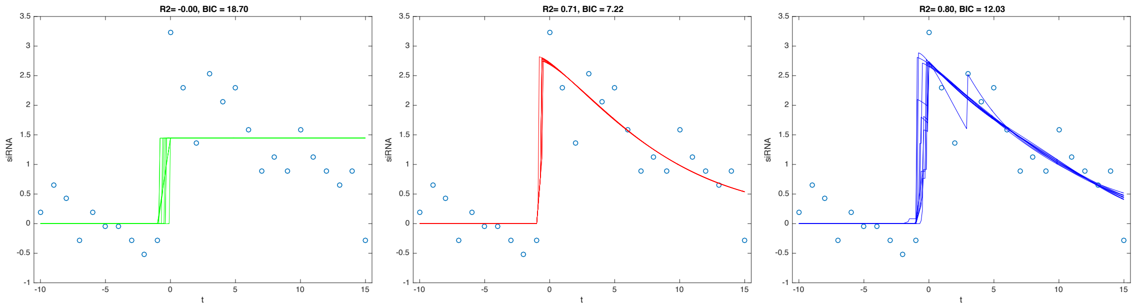


Supplementary Figure 14: The three different trajectory types displayed. For each type the ten lines display the different fit found. The dots show observations.

#### 2 eGFP response function

In order to examine the effect of the cytosolic siRNA quantity on eGFP knockdown in single cells, an explicit function for the relation is needed. The function we used is the following:

$$f(t; R, \boldsymbol{\theta}) = \exp\left(-\theta_1 R^{\theta_2} \int_0^t \exp\left(-\frac{(s-\theta_3)^2}{\theta_4^2}\right) ds\right).$$

Here, $t$ denotes time after the release event (occurring at time 0) and $R$ is the siRNA fluorescence intensity. The reason for using an exponential is the function is non-negative of the eGFP. The actual function inside the exponential is easier to understand in terms of its derivative $-\theta_1 R^{\theta_2} \exp\left(-\frac{(s-\theta_3)^2}{\theta_4^2}\right)$. Here the derivative takes it minimum at time point $\theta_3$ (after the time of release) where the value is $\theta_1 R^{\theta_2}$ (this function was found after numerical fitting). At the time of the release (time is zero) the value of the function is zero and also as time goes to infinity. The parameter $\theta_4$ controls the scale on which the derivative changes.

#### References

[1] Gideon Schwarz. Estimating the Dimension of a Model. *The Annals of Statistics*, 6(2):461 – 464, 1978.

### Supplementary Note 2 – Apparent eGFP expression increase during mitosis

We noticed an apparent increase in eGFP fluorescence short after the release event for both siGFP sequences and siLuc (Supplementary Fig. 8a). This eGFP fluorescence increase was not seen in flow cytometry data (not shown), individual cell-traces or in a previous spinning-disk based expression monitoring[8]. We did however see substantial variation of eGFP fluorescence over the cell-cycle in unperturbed cells; a gradual increase in eGFP fluorescence prior to mitosis followed by a rapid substantial drop in fluorescence after the mitotic event (Supplementary Fig. 8b). Conceivably, this is due to the rounded-up morphology during mitosis and the open pin-hole confocal setup used for the eGFP channel (to minimize the required light doses) resulting in an apparent higher signal in the large rounded up cell prior to mitosis, and a subsequent fluorescence drop upon flattening of the post-mitotic cell. This has limited effects when looking at cell ensembles were this variability averages out. However, due to the rounded up shape of the cell during mitosis, any cytosolic release events happening around mitosis will be difficult to detect, as evidenced by a lack of mitotic events for approximately 10 frames before and after a detected siRNA release event (Supplementary Fig. 8c). To correct for this effect, all cytosolic siRNA measurements were normalized to the mean eGFP fluorescence intensity in AF647-siLuc control experiments, in a time-dependent manner (Supplementary Fig. 8d).
